# Testing and training geometrical knowledge in baboons through a conceptual categorization task

**DOI:** 10.64898/2026.11.06.764828

**Authors:** Thomas Dighiero-Brecht, Lorenzo Ciccione, Mathias Sablé-Meyer, Joël Fagot, Nicolas Claidière, Stanislas Dehaene

## Abstract

Sensitivity to geometric regularity has been proposed as a signature of human cognitive singularity. Baboons can detect an intruder shape among quadrilaterals but, unlike humans, show no advantage for regular shapes such as squares and rectangles, suggesting they are not sensitive to geometric regularity. Here we asked whether, with symbolic training, baboons may show an enhanced attention to geometric features and their compositions. Using a symbolic match-to-sample task with frequent introduction of new generalization stimuli, we trained baboons and human controls to categorize stimuli as representing a given concept (A) or not (not-A) across two non-geometric domains (image category and line color) and three geometric domains (line perpendicularity, parallelism, and length equality). Baboons learned to perform above chance with all categories and generalize them to novel positive and negative exemplars (except for length equality). These findings provide evidence that baboons can extract geometric properties and associate them to arbitrary symbols. Crucially, this explicit geometry training did not result in the emergence of human-like performance in the quadrilateral intruder task: overall performance improved uniformly across shapes, but the geometric regularity effect failed to emerge. Thus, in baboons, recognizing individual geometric properties is insufficient to trigger the compositional perception of shapes that characterizes human cognition.

## Introduction

Although mathematics ranks among the most sophisticated cultural products of the human mind^1^, it appears to be rooted in proto-mathematical intuitions which are already present in the absence of any formal instruction^2−7^ or visual experience^8,9^ and even in populations without mathematical education^10−1^^4^. Schooling refines these intuitions, for instance by mapping quantities onto symbols^15^, but the capacity to extract abstract numerical and geometric properties from the environment appears to be a universal human predisposition.

To what extent is this capacity shared with other animals^16^? The approximate number sense is documented across the animal kingdom, from primates to insects^17−2^^2^; the phylogenetic roots of geometric intuitions, on the other hand, remain far less explored^23^. Moreover, the ability to compose such properties into abstract representations is considered a hallmark of human cognition^24,25^, and evidence for it in non-human primates remains debated. Negation provides a test case, since understanding “not A” requires combining an operator with its argument. Baboons learned to associate an explicit visual cue with the negation of a concept^26^, but failed to generalize this rule to novel colors and shapes^27^. In a recent study, rhesus macaques saw a cue image followed by two choice images, one identical to the cue and one different^28^. They were rewarded for choosing the identical image, unless the cue was accompanied by an abstract “negation” symboI, in which case they had to choose the different one. Because images were drawn from several hundred pictures, the correct responses could not be memorized. The monkeys first appeared to succeed, but only by treating any additional symbol as a signal to choose the different image: when a distinct “positive” symboI was added to matching triaIs, performance feII to chance. They eventuaIIy learned to use the identity of the symbols, but only after extensive training with a modified procedure. Similarly, macaques learn embedded structures and abstract sequence rules only after massive exposure, and fail to transfer them across perceptual domains^29,30^. Non-human primates may therefore possess some capacity to combine elements, but it is slow to acquire, fragile, and frequently supplanted by simpler perceptual or associative heuristics.

Geometry offers a particularly clear case. In an intruder task requiring the detection of a deviant quadrilateral, humans make fewer errors when the reference shapes are more regular: an intruder stands out more readily among squares than among trapezoids. Baboons can perform the task (although with a much larger error rate compared to humans) but show no such effect^31^. Even after thousands of rewarded trials, their performance remains similar when detecting a deviant among squares, trapezoids or random quadrilaterals, and, in the original analysis, it was mostly explained by the perceptual image properties extracted by a convolutional neural network (CNN), rather than by the conceptual or symbolic geometric properties required to model human performance. Conversely, a trial-level Bayesian reanalysis of the original baboon data reported a reliable contribution of the symbolic model, comparable to that observed in preschoolers. It proposed that much of what is labeled “symboIic” refIects rotation-invariant shape coding shared across primates^32^. Whether baboons possess a weak symbolic geometric component therefore remains debated. Neither analysis, however, indicates that baboons exhibit the human regularity effect itself. This geometric regularity effect has therefore been proposed as a signature of human singularity^24^. Although carrion crows were recently reported to succeed in a similar task^33^, they were tested with shapes deviating much more than those used in humans and primates (75% vs. 30% vertex displacement). A re-analysis of crows’ performance further indicated that it was better captured by a low-level perceptual model than by a symbolic geometric one, and correlated with baboon rather than human behavior^34^.

Nevertheless, the absence of a regularity effect in non-human animals might reflect a failure to attend to the individual properties that define it, rather than a constitutive inability to represent regularity. A square is more regular than a trapezoid because it simultaneously possesses four equal sides, four right angles and two pairs of parallel sides. A rectangle shares some of these properties but lacks equal sides. Through formal education^13^ and immersion in a highly geometrized, human-made environment^35^, humans learn to attend to the presence and absence of such properties and to combine them. Does the failure of non-human primates therefore reflect an absolute cognitive boundary, or merely a failure to attend to the relevant properties?

To address this question, we decomposed geometry into its elementary properties, and taught them to baboons separately. Using a previously used symbolic match-to-sample categorization task^36^, we trained baboons and human controls to categorize stimuli as belonging to a concept (A) or not (not-A). We asked whether baboons could learn naturalistic categories and their negation^26^ and whether they could extract perpendicularity, parallelism and length equality (and their absence), as well as color as a non-geometric control, and generalize them to novel exemplars independently of irrelevant dimensions. Crucially, we administered the quadrilateral intruder task before and after this curricular training, to test whether learning individual geometric properties suffices to produce a human-like regularity effect. We hypothesized that it would not: even if individual properties can be learned, possibly by relying on feature detectors in the ventral visual stream^37^, the intruder task requires their simultaneous evaluation and integration across a whole shape, so if baboons lack the compositional abilities needed to combine them, isolated-feature training should be insufficient to change their behavior in this task.

## Results

### Both humans and baboons learn to categorize naturalistic images

Each participant completed five blocks (naturalistic images, perpendicularity, parallelism, length equality and color) in a randomized order. On each trial, a stimulus appeared on screen, followed by two abstract symbols serving as response buttons; participants had to seIect the symboI corresponding to the stimuIus’s category (A or not-A) and received feedback (Figure 1A; see Methods). Each block used its own two symbols, drawn at random for each participant from a set of ten, with their positive and negative meanings aIso randomIy assigned. Hence, the symboIs used to respond “crow” and “not crow”, for instance, differed from those used for “right angIe” and “not right angIe”.

**Figure 1.**
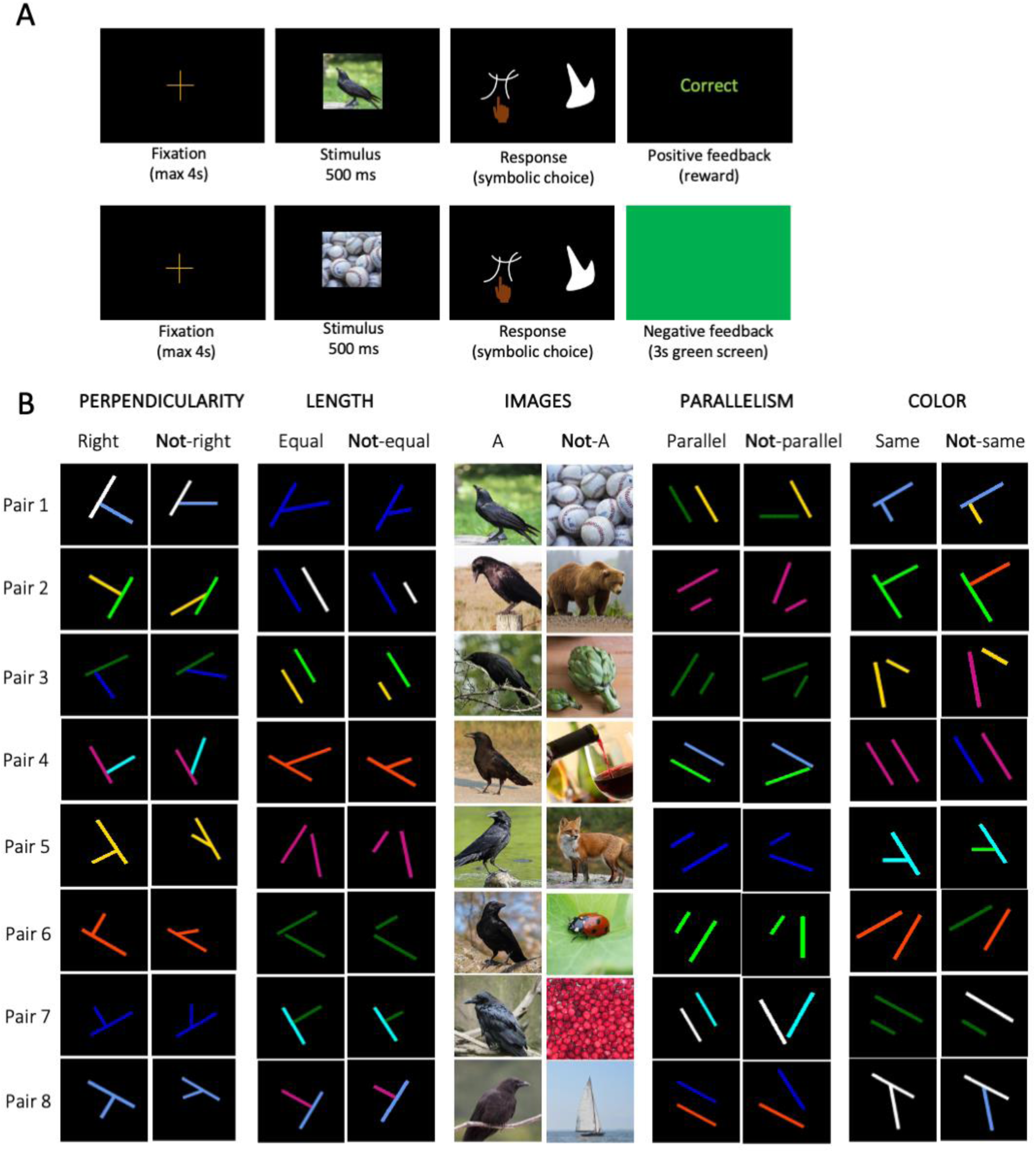
A symbolic match-to-sample task to probe conceptual categorization of geometric properties and naturalistic categories. (A) Two example trials from the naturalistic-image block. Participants start each trial by touching (baboons) or clicking (humans) the fixation cross. A stimulus is shown for 500 ms, followed by two arbitrary symbols. The participant selects the symbol associated with the stimulus’s category and receives species-specific feedback. (B) Complete stimulus set for one participant. Each block tests one concept through eight successive pairs. Each pair comprises one positive stimulus (belonging to the concept) and one negative stimulus (not belonging to it), presented one at a time in random order. Participants advance to the next pair once they reach 70% accuracy on both. For naturalistic images, one category is randomly assigned as positive, and each negative stimulus is drawn from a different one of the eight remaining categories.

We begin by reporting performance with naturalistic images, which allowed us to probe positive and negative concepts for ecological stimuli. Each participant was assigned a specific category (the “positive” stimuIi; e.g., *crows*) and, at each of eight successive training levels, one positive exemplar was paired with an exemplar drawn from one of the eight remaining categories (the “negative” stimuIi; e.g., *ladybugs*). A new pair was introduced only once participants reached 70% correct responses in that training level (Figure 1; see Methods). Note that, throughout this paper, the term *pair* should not be interpreted as the simultaneous on-screen presentation of two stimuli; it will simply refer to the coupling between one positive and one negative stimulus category, used to define a single training level.

#### Learning rate

Both species showed clear evidence of learning (Figure 2A). Baboons required progressively fewer trials to reach the accuracy threshold across pairs, plateauing at roughly 200 trials from pair 5 onwards (exponential decay fit to pairs 2−8, R² = 0.797), while humans rapidly approached ceiling performance, i.e., 16 trials, reaching it by pair 4.

**Figure 2.**
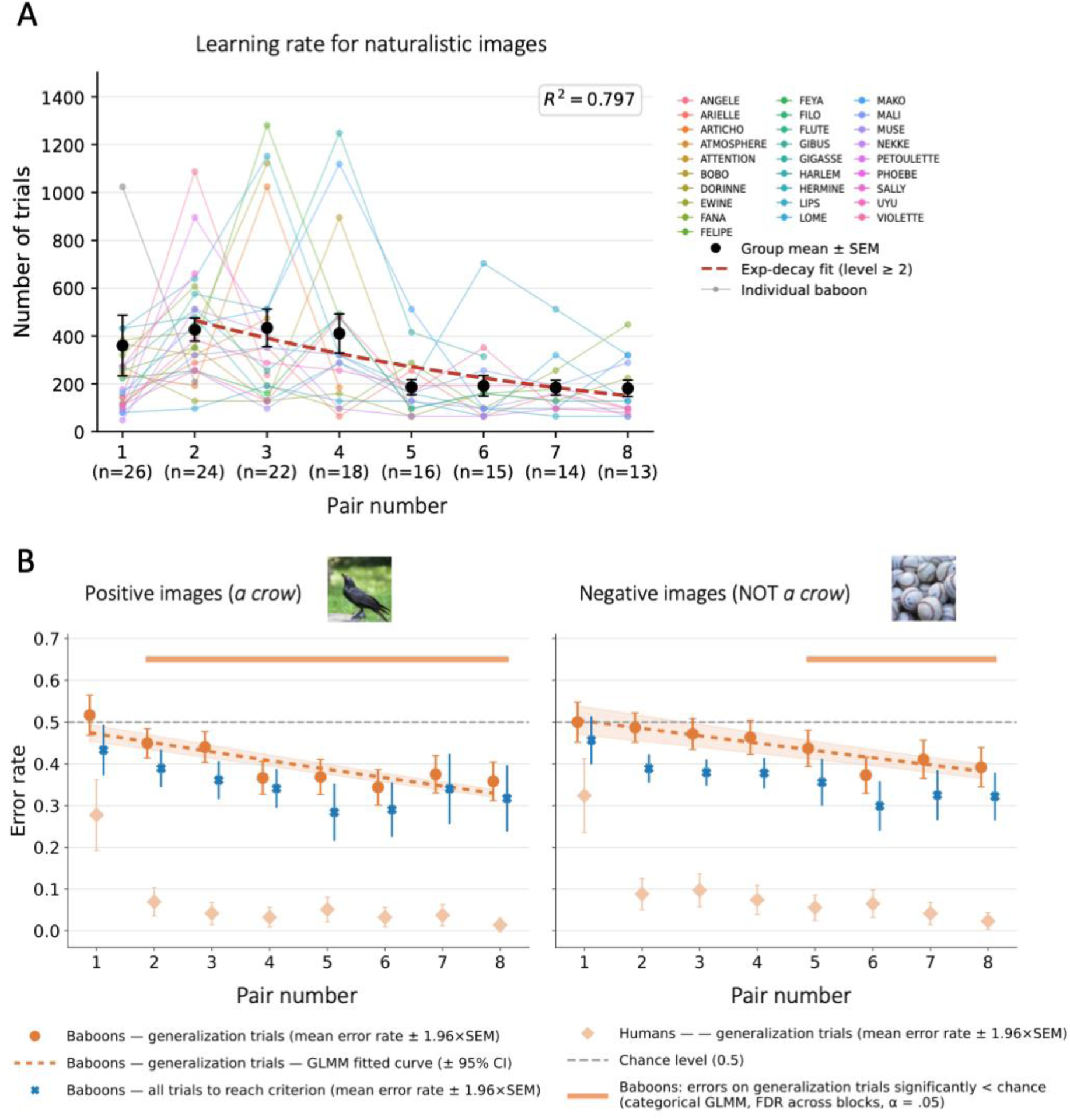
Baboons can categorize ‘A’ and ‘non-A’ stimuli through implicit negation. (A) Learning rate: number of trials to reach the 70% criterion at each training level. Colored lines show individual baboons; black dots show the group mean ± SEM. The red dashed curve is an exponential decay fitted to group means at levels 2−8 (R² = 0.797). The number of baboons per level is given on the x-axis. (B) Error rate on generalization trials (first batch of each training level) for positive trials (left; exemplars of the assigned category, e.g., crows) and negative trials (right; exemplars of any other category). Symbols are as follows: • Orange circles: baboon mean error rate on generalization trials ± 95% CI (1.96 × SEM, computed across trials pooled over animals). • Orange dashed line: GLMM fit ± 95% credible interval. • Light orange diamonds: human mean error rate ± 95% CI (1.96 × SEM across participants). • Blue crosses: baboon mean error rate across all trials to criterion ± 95% CI (1.96 × SEM across animals). • Grey dashed line: chance. • Orange bars along the x-axis: levels with below-chance error (categorical GLMM, Benjamini− Hochberg FDR across 80 contrasts, a =.05).

#### Error rate on generalization trials

To assess whether participants applied the acquired rule to novel exemplars, independently of within-pair learning, we examined the error rate on the first batch of 64 trials encountered at each new training level (hereafter, generalization trials), separately for positive and negative stimuli (Figure 2B). For each block, we fitted 2 different mixed-effects logistic regression, one continuous and one categorical (see Methods).

In baboons, generalization error decreased across pairs for both stimulus types (positive: β = −0.087, z = −17.49; negative: β = −0.071, z = −8.29; both p <.001; TabIe S1), but reliable generalization emerged later for negative stimuli: the error rate was significantly below chance from pair 2 onwards for positive stimuli, but only from pair 5 onwards for negative stimuli (FDR-adjusted p <.05; Table S2). At pair 8, both error rates were significantly below chance, but remained higher for negative (38.3%) than for positive stimuli (33.0%; difference: z = 7.55, p <.001). The difference in onset, pair 2 versus pair 5, likely quantifies the cognitive cost of implicit negation: baboons required approximately three additional training levels before their rejection of negative stimuli departed reliably from chance. Human participants generalized both categories after only a few pairs, with error rates reaching approximately 5% by pair 3 for positive images and by pair 5 for negative images.

#### Immediate generalization

Below-chance error on generalization trials could arise in two ways. Baboons might immediately apply the acquired rule to the new pair (immediate generalization), or they might still need to learn the new pair, but do so faster than the previous ones because of prior training (faster learning). To separate these two possibilities, we split the generalization batch of the image block into four consecutive quarters of 8 trials per stimulus type, in the 13 baboons that reached pair 8 (Figure S1). For positive stimuli, error was already below chance in the first quarter (38.2%, p_FDR_ <.001) and remained so throughout. For negative stimuli, error was at chance in the first quarter (51.4%, p_FDR_ =.82) and fell below chance only from the second quarter onwards (42.9%, p_FDR_ <.001). The same pattern held when the analysis was restricted to pairs 5-8, where negative generalization over the whole batch was reliable (first quarter: 50.2%, p_FDR_ =.61; Figure S1B). Baboons therefore did not immediately reject a new non-A category; they only did so by learning from feedback within the batch. Such learning, however, became faster from pair to pair: the number of trials to criterion decreased across pairs (Figure 2A), and so did the error rate on negative generalization trials (Table S1). Negation thus showed savings due to faster learning, but not to immediate generalization.

### Baboons learn to categorize geometric properties

The four remaining blocks used stimuli made of two-line segments. The color block, in which the concept to be learned was whether the two lines shared the same color or not, served as a non-geometric perceptual control: baboons learned and generalized this non-geometric rule, and negative stimuli eventually reached lower error rates than positive ones (pair 8: 27.9% versus 42.6%; Supplementary Results; Figure S2). The three geometric blocks probed the angle that the two lines form (perpendicular or not), their parallelism (parallel or not), and their relative length (same length or not).

#### Learning rate

For baboons, all three geometric blocks showed a non-monotone learning rate profile, with a steep decline from pair 3 onwards, stabilizing at approximately 150−300 trials by pairs 7−8 (exponential decay fit to pairs 2−8: angles, R² = 0.997; parallelism, R² = 0.952; length, R² = 0.991; Figure 3, left panels). Human participants learned all three rules with trial counts close to optimal performance throughout the task.

**Figure 3:**
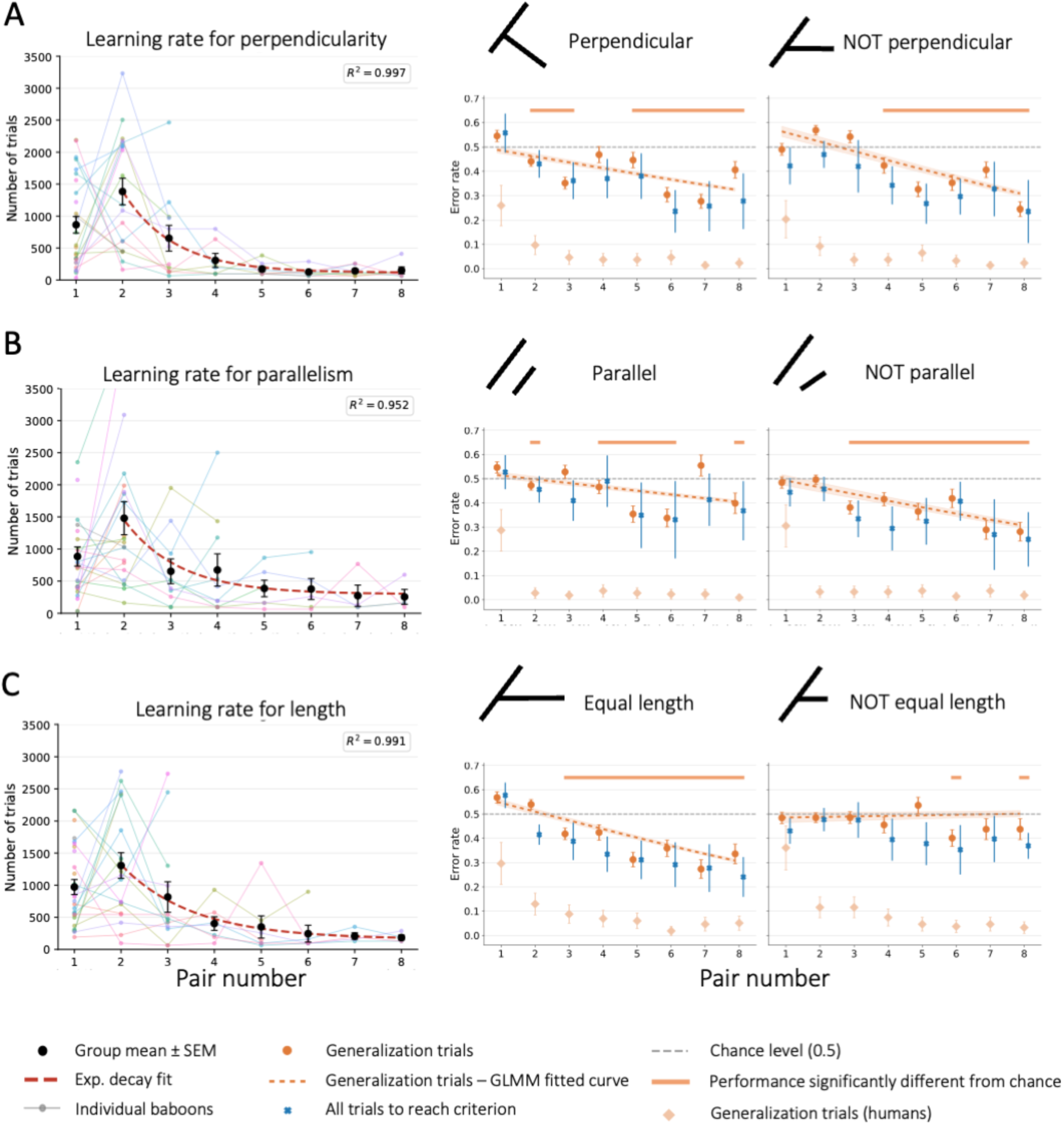
Baboons learn and generalize elementary geometric properties and, except for length, their negation. Perpendicularity (A), parallelism (B) and length equality (C). Left panel: learning rate, as in Fig. 2A (exponential fits: angles, R² = 0.997; parallelism, R² = 0.952; length, R² = 0.991). Right panel: error rate on generalization trials, as in Fig. 2B. The left subpanels show positive trials (perpendicular, parallel, equal-length) and the right subpanels show negative trials (non-perpendicular, non-parallel, unequal-length).

#### Error rate on generalization trials

Generalization of the positive concept was successful in all three blocks (Figure 3, right panels): the error rate was significantly below chance from pair 2 onwards for perpendicularity and parallelism, and from pair 3 onwards for length, although with slight oscillations in the middle pairs (Table S2), and remained so until pair 8 (angIes: 32.5%, z = −50.69; paraIIeIism: 40.6%, z = −25.83; Iength: 30.8%, z = −54.91; aII p <.001; Table S1). For negative stimuli, the pattern depended on the geometric property. For perpendicularity and parallelism, negative generalization was robust (below chance from pair 4 and pair 3 onwards, respectively; pair 8: 30.9% in both blocks, p <.001) and ultimately more accurate than positive generaIization (difference at pair 8: angIes, z = −3.62; parallelism, z = −20.05; both p <.001), a reversal of the asymmetry observed for naturalistic images. Such reversal was observed for color as well (see above). For length, by contrast, negative generalization was weak and late. When all animals were pooled, the error rate on unequal-Iength stimuIi did not decrease significantIy across pairs (β = +0.009, z = 1.81, p =.071) and was at chance at pair 8 under the continuous model (50.1%, z = 0.23, p =.82); the categorical model detected only small below-chance deviations at pairs 6 and 8 (48.5% at pair 8, FDR-adjusted p =.013). In either case, performance remained far from that reached on equal-length stimuli (trial-type × training-IeveI interaction: β = +0.15, z = 37.55, p <.001). Human participants showed near-floor error rates from pair 2 onwards for both trial types in all three blocks, with no positive-negative asymmetry.

Because animals that failed to progress stopped contributing data, and attrition was partly performance-based (completers made slightly fewer errors at pair 1 in the geometric blocks; Figures S3-S4, Table S4), we refitted all models to the animals that reached pair 8 in each block (n = 4-13). In this other sample, above-chance generalization to positive stimuli and the positive−negative asymmetry were preserved in all five blocks, although the trial-type × training-level interaction was no longer significant for images and angles, and completers of the length block also generalized to negative stimuli by pair 8 (SI, *Robustness to attrition*).

To summarize, baboons learned and generalized all three geometric categories, as well as the implicit negation of perpendicularity and parallelism, whereas generalization of the negation of length equality remained, at best, marginal.

### Training on basic geometric properties does not produce a human-like sensitivity to geometric regularity in baboons

The fundamental question we asked was whether prolonged exposure to geometric stimuli, and learning to categorize their basic properties using specific symbols, would produce a qualitative shift in sensitivity to geometric regularities in quadrilaterals. To answer this question, we tested the baboons before and after the geometry training on the intruder task designed by Sablé-Meyer et al^31^. Baboons’ performance before training aImost exactIy replicates the starting point from the original study (Fig 4A pink versus blue; see the Methods for further details). At the end of training, the apparent superiority of the originally reported data (purple line) reflected the additional practice on the oddball task itself that they included. Error rate also fell after the geometry training on our categorization task (Figure 4A, green line), and did so in every individual tested across all three sessions. Crucially, this improvement was uniform across shapes: the error advantage for regular over irregular quadrilaterals did not grow across sessions, so no human-like regularity effect emerged (see Figure 4A, grey dotted line for human data from our original study).

**Figure 4.**
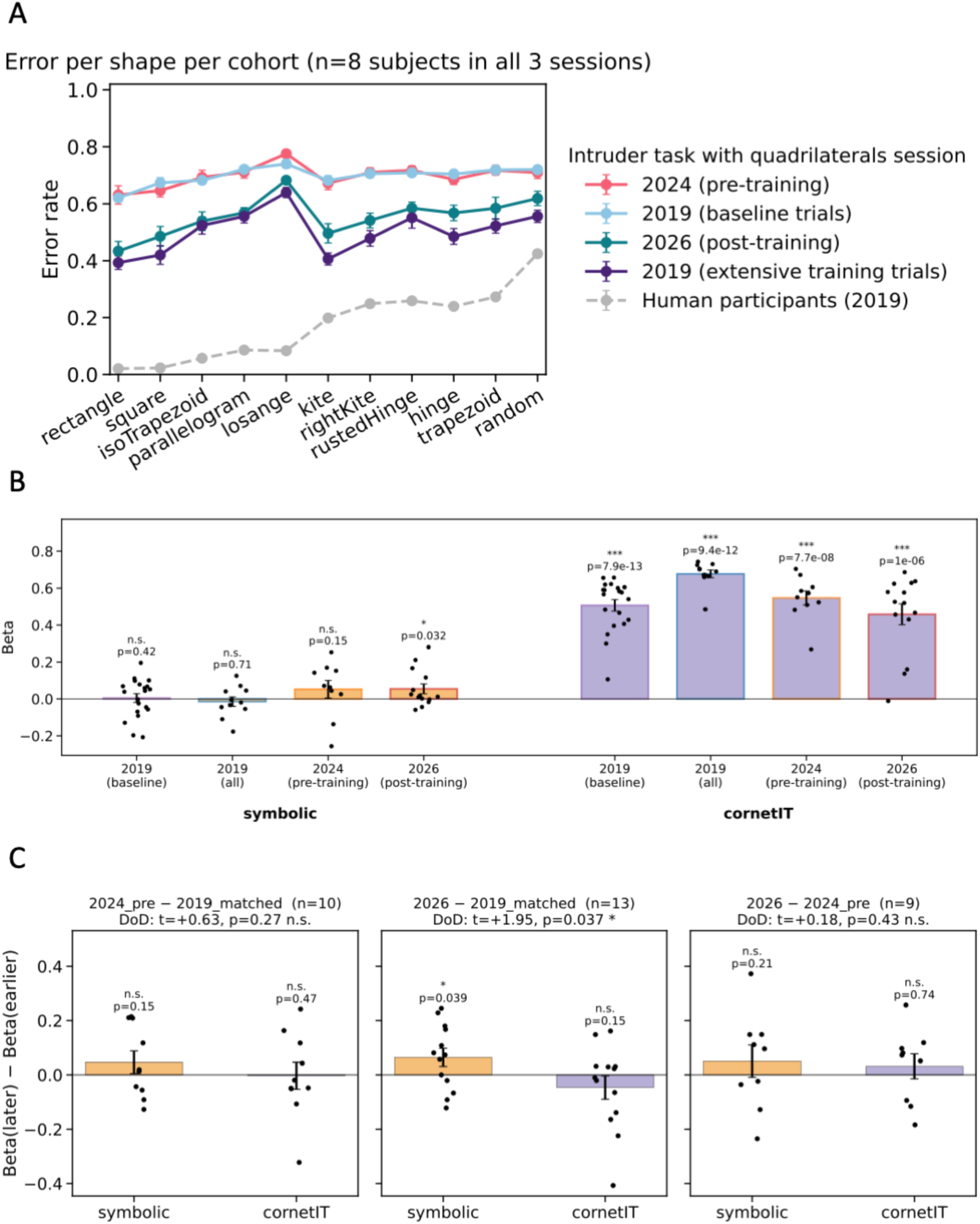
**Geometric training does not produce a human-like regularity effect in baboons.**“2019 (all)” denotes the original data as reported in ref. 31. “2019 (baseline)” is the same data restricted to the first 30 bIocks (≈2,640 triaIs) per animaI. “2024 (pre-training)” and “2026 (post-training)” are Iikewise restricted to the first 30 bIocks, or all trials for animals with fewer. • (A) Mean error rate (± SEM across animals) per quadrilateral, for the eight baboons tested in all three sessions. Shapes are ordered as in ref. 31. Data from human participants (experiment 1 from ref. 31) are shown in dashed grey line • (B) Coefficients from the per-animal, per-session regression error ∼ symbolic + CorNet-IT. Predictors are z-scored so that higher values indicate conditions predicted to be harder. Bars show the mean across animals, dots show individual animals, and error bars show SEM. Tests are one-sided one-sample t-tests against zero (n.s., not significant; *p < 0.05; **p < 0.01; ***p < 0.001). • (C) Within-animal change in coefficients between sessions, restricted to animals tested in both sessions (n in panel titles). Tests are one-sided paired t-tests (symbolic: increase; CorNet-IT: decrease). Panel titles also report the difference-of-differences test between terms.

Following our previous work^31^, we also regressed error rate, computed per condition, on the predictions of a symbolic model and of a convolutional neural network (CorNet-IT; Figure 4B). Under this condition-level analysis, baboon performance was again far better captured by CorNet-IT than by the symbolic model. The symbolic model did show a very small but significant influence after our training on geometric properties (p =.032, Figure 4B). However, this increase occurred at least partially prior to said training: there was a small but significant difference between the new data with and the trial-matched original data (p = 0.039; Figure 4C, 2026 − 2019_matched), but it did not increase between the pre-training replication and the post-training session (p = 0.21, n.s.; Figure 4C, 2026 − 2024_pre). Furthermore, caution is needed because these effects were small, uncorrected, and there was no corresponding decrease in the influence of the convolutional network model (Figure 4B). Although a modest shift toward the symbolic model may have emerged over the years, it cannot be attributed specifically to our curriculum, but rather to the animaIs’ cumuIative experience with geometric material, which includes repeated testing on the intruder task and other geometric and compositional tasks. At the individual level, no animal showed a significant symbolic contribution after training once corrected for multiple comparisons (Table S6).

Because the animals varied widely in how far they progressed through the curriculum, no animal completed all three geometric blocks, and 17 of the 28 completed none (Table S5), we also asked whether the amount of geometry training predicted the change in intruder-task performance. Among the nine animals tested both before and after training, neither the number of geometric stimulus pairs encountered (5 to 19) nor the number of geometric blocks completed (0 to 2) correlated with the reduction in error rate, with the change in the symbolic coefficient, or with the change in the CorNet-IT coefficient (aII |p| ≤.27, aII p ≥.48; Figure S5, Table S6), including when controlling for baseline performance, total number of trials and age (Supplementary Results). The animal with the least geometry training (5 pairs, no block completed) improved as much as the most-trained animals (error-rate reduction of 18 percentage points, versus 7−21 points). However, nothing in these data suggests that the size of the improvement depended on the amount of geometry training rather than on practice with the task.

## Discussion

This study provides three main findings. First, baboons can learn elementary geometric properties (parallelism, perpendicularity and length equality), bind them to arbitrary symbols and generalize them to novel exemplars. These results provide evidence of proto-mathematical abilities in non-human primates beyond the approximate number sense. Second, with naturalistic images, baboons generalized a negative category to entirely novel exemplars, a hint that they can represent “not-A,” although, as in previous studies, the evidence for a genuine negation operator remains limited. Third, acquisition of this geometric lexicon did not suffice to trigger a strong change in how the animals viewed quadrilaterals: extensive training did not produce a human-like sensitivity to geometric regularity. This suggests that what baboons lacked was either the compositional machinery that combines such primitives, or sufficiently abstract representations of whole shapes. We discuss these two possibilities below.

### Proto-geometric abilities in baboons

The extraction of geometric features has been far less studied in animals than numerical abilities^23^. Here, baboons learned to categorize parallelism, perpendicularity, and length equality. They generalized these categories to novel exemplars despite variations in orientation and absolute line length that ruled out reliance on a fixed image template, although only a subset of animals completed all eight training levels and learning rates varied widely (SI Appendix). These abilities concern the very primitives that, in humans, compose the representation of geometric shapes^24,38,39^. These results extend the known proto-mathematical repertoire of non-human primates from number^17^ to geometry. They also show that baboons’ previously reported insensitivity to geometric regularity^31^ cannot be attributed to an inability to perceive its constituent properties.

### A hint of implicit negation

#### The case of naturalistic images

Previous attempts to demonstrate compositional negation in baboons have been equivocal. Baboons learned to associate an explicit visual cue with the negation of a concept^26^, but did not generalize this rule to novel colors and shapes^27^. Our paradigm replaced the explicit inversion cue with an implicit negation: one arbitrary symbol for a naturalistic category, another for its negation. Under these conditions, baboons generalized beyond trained exemplars. Because each new negative stimulus belonged to a category that had never been seen before, providing correct “not-A” responses required recognizing the absence of category “A”. This behavior is functionally equivalent to set complementation. It also differs in an important way from the macaque negation task described in the Introduction^28^. In that task, the image to be negated was always visible on the screen, and one of the two response options was an exact copy of it. Negation therefore amounted to choosing the option that differed from a visible image, a conditional same/different rule of a kind that monkeys have long been known to master. In our task, by contrast, a single image was shown and no exemplar of the trained category was present on negative trials. Responding “not-A” required comparing a novel image with a category held in memory. Moreover, both species generalized the positive category earlier than the negative one (pair 2 versus pair 5 in baboons), paralleling the well-known cost of negation in humans^40^. The much larger learning period in baboons than in humans is consistent with the proposal that language and compositional thought^24,25^ amplify the efficiency with which humans construct such categories.

Two caveats nonetheless prevent us from concluding that baboons used a negation operator. First, baboons did not reject new non-A exemplars from the start. During the first 8 trials of each new pair, error on negative stimuli was at chance, even at pairs 5-8, and it fell below chance only after feedback (SI Appendix, Fig. S1). Their progress on negation thus reflects savings, meaning faster learning of each new non-A pair, rather than immediate application of a rule. Second, the asymmetry may partly reflect familiarity, since new positive exemplars resembled previously trained ones whereas new negative exemplars did not. This difference may also explain why the macaques, once trained, applied their negation symbol to new images on every trial, whereas our baboons did not immediately apply their negative symbol to new categories. Negating a category stored in memory may be harder than rejecting a copy of an image that is directly visible.

#### Perceptual accounts of the reversed asymmetry for geometric stimuli

Across the three geometric blocks and the non-geometric color block, the positive-negative asymmetry or “negation cost” was not replicated in the same direction as with naturalistic images. Instead, the behavioral difference between positive and negative stimuli took qualitatively different forms depending on the dimension under test, suggesting that for artificial and geometric stimuli, what we initially envisaged as “negation learning” might in fact reflect other cognitive strategies.

For the parallelism and perpendicularity blocks, remarkably, negative generalization was faster and ultimately more accurate than positive generalization. Error rates on negative trials dropped earlier and reached lower asymptotic values, a reversal of the pattern observed with naturalistic images. Thus, it was easier for animals to label two lines as *not* parallel or *not* perpendicular, than to decide that they were. For the color block, a similar reversal emerged in the later pairs, with negative stimuli eventually reaching lower error rates than positive stimuli despite a slower initial learning trajectory. We propose that this reversal can be explained at two related levels.

At the level of neural representations, Pramod and Arun^41^ showed that in monkey inferotemporal cortex, the region critical for object recognition, the neural vectors for stimuli composed of two identical mirror-image parts (i.e., symmetric objects) are paradoxically more *distinctive* from each other than those for stimuli composed of non-matched parts (asymmetric objects). This is because, in high-dimensional vector space, the addition of the vectors associated with each feature sends symmetric objects farther away from the mean than asymmetric objects. This simple property, which readily explains the faster perception or “pop-out” of symmetricaI objects, may aIso expIain the baboons’ performance with Iine stimuli, at least in the parallel and color blocks. It predicts that negative exemplars (composed of parts that differ from one another) should cluster more tightly in IT representational space, forming a more coherent, easily learnable category, while positive exemplars (each composed of two matching parts) should be more individually distinctive and therefore harder to group into a single category. This interpretation may also apply to our right-angle stimuli, once one notices (as Euclid did) that a right angle splits a line into two equal angles, and therefore our positive right-angle stimuli exhibited a symmetry that their negative counterparts did not. More generally, such a representational geometry could provide the mechanism by which the relevant features are detected in the first place, yielding a feature-level code on which downstream classification can operate.

At the level of individual stimuli, two lines of different color, two non-parallel lines, or two lines subtending a non-right angle make for a visual display which is more heterogeneous than their positive counterparts. A rich body of prior work on same−different discrimination in baboons has established that these animals are exquisitely sensitive to visual entropy and readily exploit it as a classificatory cue^42,43^. Similarly, pigeons learn faster when the heterogeneous rather than the homogeneous displays are the reinforced category, both with 16-icon arrays of high versus low entropy^44^ and with two-item pairs of colored shapes^45^. Entropy, however, is only defined relative to a given encoding of the stimulus. Computed over pixels, our positive and negative stimuli have similar entropies; they differ only once each display is described in terms of the color, orientation or angle of its parts, which are the very properties under test. Entropy therefore describes the reversal rather than explaining it independently, and would presuppose a feature code such as the one described above.

These patterns may be also better explained by recoding the negative category as a distinct positive category, than by conceptual negation. Non-parallel lines, for instance, are lines that would eventually cross if prolonged. No such recoding was possible for naturalistic images, whose negative exemplars (e.g., a bear, a fox, a boat) shared no common feature. The length block fits this account. Unlike non-parallel, non-perpendicular or differently colored lines, unequal-length lines were not consistently secant, chromatically heterogeneous or asymmetric at their junction, and thus offered no positive cue, and negative generalization remained marginal at best. Since equal-length lines are also made of matched parts, representational geometry alone would have predicted a reversal here too; its absence suggests that recoding drives the advantage of negative stimuli. Such reliance on the simplest available cue has a parallel in the macaque negation study^28^. There, the monkeys initially chose the non-matching image whenever any additional symbol appeared next to the cue, regardIess of which symboI it was, and faiIed once a second, “positive” symboI was introduced. They learned to use the identity of the symbols only after 20,000 to 80,000 further trials. Even then, the authors acknowledge that this success may reflect an association between each symbol and a response strategy (choose same versus choose different), rather than a symbolic negation operator. Take together with the naturalistic-image results, our findings strengthen, but do not settle, the case for implicit negation in baboons.

### Geometric lexicon without syntax: lack of emergence of a geometric regularity effect

Our design allowed a direct test of whether the absence of a geometric regularity effect in baboons^31^ reflects a failure to attend to the constituent properties of regular shapes. Our results do not support this explanation. After training on parallelism, perpendicularity and length equality, baboons did not develop the human pattern of making fewer errors on more regular shapes, such as squares and rectangles. Their overall error rate decreased after training, but uniformly across shapes, and by an amount unrelated to how much geometric feature training each animal had received. Furthermore, the results of the baboons were still massively captured by a convolutional neural network. Still, the relative contribution of the symbolic model increased slightly over the years. This increase was not specifically tied to our curriculum, but it suggests that cumulative experience with geometric material may slightly strengthen a symbolic component that is present but weak.

Two other lines of work point in the same direction. A small but significant language-of-thought contribution, much weaker than in humans, emerged when baboons were forced to slow down their responses and were tested in a delayed match-to-sample task with a broad set of shapes^46^. In addition, a trial-level Bayesian reanalysis of the original intruder data reported a reliable symbolic contribution in baboons^32^. That result departs from our condition-level analysis and may reflect differences in analytic resolution, since averaging across trials can mask weak but systematic effects. Rudimentary symbolic geometric representations may thus be partially available to baboons. None of these analyses, however, shows that baboons exhibit the regularity effect itself, the systematic advantage for regular shapes that characterizes human performance^31^ and that our training failed to induce.

Our findings provide, to our knowledge, the most direct evidence so far that human sensitivity to geometric regularity goes beyond the successful learning of elementary geometric properties. Perceiving a square as more regular than a trapezoid requires knowing what parallelism, perpendicularity and equal lengths are, but it also requires evaluating their co-presence across a single shape and deriving a holistic judgment from their combination. Two accounts could explain why baboons failed at this step.

On the language-of-thought account, regular shapes are simpler because they can be described by shorter programs in a recursive, compositional mental language^24,38,39^. Baboons would have acquired part of the lexicon of this language but would lack the syntactic machinery that combines its primitives into compact descriptions.

On a continuity account, the human advantage instead reflects more powerful analog, rotation-invariant representations of whole shapes, for which the symbolic feature model is partly a proxy. In both humans and monkeys, symbolic-model weights correlate with those of a rotation-invariant version of a CNN and increase when rotated shapes must be matched^32^. Baboons would then lack not a syntax, but sufficiently abstract shape codes, which training on isolated line pairs would not supply.

The present data cannot fully separate these accounts, but two observations suggest that rotation invariance alone may not suffice. First, our geometric stimuli varied in global orientation, so baboons learned to recognize perpendicularity and parallelism across orientations, yet this did not translate into a regularity effect. Second, the symbolic model used here and in Li et al., 2026^32^ scores shapes by their individual Euclidean properties. It therefore indexes the geometric lexicon, which baboons can demonstrably acquire, rather than the compression of these properties into a compact description, which is what the regularity effect requires. Distinguishing the two accounts will require models that explicitly contrast program length with rotation-invariant holistic similarity, applied at the trial level to both species.

A promising direction for future work would therefore be to explicitly teach the compositional structure itself. For instance, baboons could be trained to associate a symbol with a conjunction of learned properties (e.g., lines that are both parallel and of equal length) and tested on novel shapes. Another candidate is the geometric tracing task used by Tian et al., 2026^47^. Conversely, if the continuity account is correct, training that promotes orientation-invariant recognition of whole shapes should be more effective in inducing a regularity effect than training on combinations of properties. Contrasting these two forms of training would help determine whether, and how, the gap between humans and non-human primates that our training on individual geometric features could not close can be bridged.

## Methods

### Participants

#### Baboons

A group of 28 Papio papio baboons (21 females, 7 males) with ages ranging from 2.1 to 29.7 years (mean 16.7 years), were tested at the CNRS primate facility situated in Rousset-sur-Arc, France. The baboons lived in social groups with free access to 14 automated learning devices, each featuring a touch screen, a food dispenser, and a radio-frequency identification (RFID) reader that identified each animal and allowed it to resume the task where it had left off at each new entrance (SI Methods). The experiment was approved by the French Ministry of National Education (approval no. APAFIS-2717-2015111708173794-157 V3) and conformed to the relevant regulatory and ethical standards.

#### Humans

We included 27 adults (16 women, 11 men) with ages ranging from 21 to 34 years (mean 25.2 years), recruited via the PRISME platform of the Paris Brain Institute and compensated 15 euros for their participation. The experiment was approved by the Paris 8 University ethical committee (CE-P8-2024-07-01).

### Experimental procedure

#### Symbolic match-to-sample task

The task was analogous to Ciccione et al^36^ (Figure 1A). A yellow fixation cross was presented until it was touched, or for a maximum of 4s, followed by an experimental stimulus, presented for 500 ms. Two arbitrary response symbols were then displayed, one associated with the positive (A) and one with the negative category (not-A), and the participant had to touch one of the two within 4s; the position of the symbols was randomized across trials. Correct responses were rewarded with a small amount of food, whereas incorrect responses were followed by a 3-s green timeout screen. Response symbols were abstract shapes devoid of straight lines and sharp angles, assigned in maximally dissimilar pairs. We grouped the symbols into 5 pairs, each of which was randomly assigned to a designated block. Within each pair of symbols, one symbol was randomly defined as positive, the other one as negative (see SI Methods for further details).

#### Blocks and stimuli

All participants performed five blocks in a randomized order (Figure 1B), each designed to test a different concept and its implicit negation: the parallelism of two lines (versus non-parallel lines), the presence of a right angle between them (versus a non-right angle), the equality of their length (versus unequal lengths), their same color (versus different colors), and the membership in a naturalistic category (e.g., a whale, versus any other category).

Naturalistic images were drawn from 9 categories of 8 exemplars from the THINGS initiative^48^: for each participant, one category was randomly assigned as positive, and each negative exemplar was drawn from a different remaining category. Geometrical stimuli consisted of two colored straight lines generated with the *drawvsg* Python library; the two members of each pair differed only along the dimension under test, while task-irrelevant parameters (global orientation and absolute length of the lines) were varied to prevent baboons from exploiting low-level perceptual regularities (SI Methods).

#### Structure of the task

Within each block, up to eight stimulus pairs were introduced successively. At each training level, the participant was tested exclusively on one pair until reaching the success criterion of 70% correct responses on both positive and negative stimuli, evaluated over batches of 32 trials for pair 1 and 64 trials for subsequent pairs. Note that the term pair refers exclusively to the conceptual pairing of one positive and one negative stimulus: on every trial, only a single stimulus was displayed. Human participants completed an almost identical version of the task: same stimuli, batches of 8 trials (pair 1) and 16 trials (subsequent pairs), responses made by mouse click, and visual rather than food feedback.

#### Quadrilateral intruder task

Before and after the main task, baboons completed the same intruder detection task introduced by Sablé-Meyer et al.^31^, in which they had to identify, among six quadrilaterals, the one deviating from a reference shape (11 reference shapes × 4 deviant shapes; SI Methods). The pre-training (2024) and post-training (2026) sessions were compared with the data of the initial study (2019). Because performance improves with practice and the three sessions differed in length, all cross-session comparisons were restricted to the first 30 bIocks (≈2,640 triaIs) per animaI, unIess otherwise stated.

### Statistical analysis

#### Learning rate

For each block and pair, learning rate was quantified as the number of trials required to reach the success criterion, summarized across subjects as mean ± SEM. Group-level curves at pairs 2−8 were fitted with a three-parameter exponential decay, y = a·exp(−b·x) + c, which provided the best or near-best fit among four candidate functions across blocks (SI Methods).

#### Generalization across training levels

Generalization was measured on the first batch of trials of each training level (generalization trials, 16 positive and 16 negative trials for pair 1, 32 positive and 32 negative from pair 2 onwards), in which the new stimulus pair was presented for the first time. For each block, baboon errors were analyzed with a mixed-effects logistic regression (binomial GLMM, fitted by variational Bayes) with a random intercept per animal. A first model included training level (continuous, centered on pair 8), trial type (positive, negative) and their interaction. It tested whether the error rate at pair 8 differed from chance, whether it decreased across pairs, and whether positive and negative trials differed on either measure. A second model treated each training level as a separate category, making no assumption about the shape of the learning curve. In this model, we tested pairs against chance (one-sided tests; 5 blocks × 8 pairs × 2 trial types = 80 tests, Benjamini− Hochberg false discovery rate correction^49^). Fitted curves in the figures come from the first model; below-chance markers come from the second (SI Methods)

#### Robustness to attrition and training dose

Both models were refitted to the animals that completed each block, and pre-selection was assessed by comparing the pair-1 error rate of completers and non-completers. The association between the amount of geometry training and the change in intruder-task performance was assessed with Spearman and partial correlations (SI Methods).

#### Intruder task

Performance was quantified, for each animal and session, as the error rate per reference shape. Following Sablé-Meyer et al^31^, we regressed, for each animal and session, the error rate across the 44 conditions on the predictions of a symbolic model and of the IT layer of the CorNet-S convolutional neural network (error ∼ symbolic + CorNet-IT)^50^. The resulting per-animal coefficients were tested against zero within each session and compared between sessions in the animals that performed both (SI Methods).

## Acknowledgments

This research was supported by INSERM, CEA, Collège de France, Sorbonne Université, ERC grant “MathBrain” to S.D., and the *Fondation pour la Recherche Médicale* foundation, FDM program, grant number FDM202206015336 to T.D.B. This work was also supported by a grant from the Agence nationale de la recherche (ANR-16-CONV-0002 - ILCB) as well as a grant from the French government managed by the Agence nationale de la recherche as part of the France 2030 program, under the reference ANR-23-IAIIU-0010. We thank the team from the ‘‘Station de primatoIogie’’ for their heIp in conducting the study, as weII as the PRISME team from Paris Brain Institute and Polyana Araujo da Silva for her help with data collection with human participants.

## Supplementary Materials

### Supplementary Results

#### Immediate generalization and savings in the image block

On generalization trials, error rates fell below chance for both positive and negative stimuli (Figure 2B; Table S2). This can reflect two different processes. In immediate generalization, the rule is applied correctly to a new pair from its very first presentations. In savings, the new pair still has to be learned from feedback, but more quickly than earlier pairs. To separate them, we split each generalization batch into four consecutive quarters of 8 positive and 8 negative trials, in the 13 baboons that reached pair 8 (Figure S1; SI Methods).

Pooling pairs 2-8, the two stimulus types behaved differently (Figure S1A). On positive trials, error was below chance from the first quarter (0.38, z = −6.47, p_FDR_ <.001) and in every subsequent quarter (all p_FDR_ <.001), with only a slight decline over the batch (0.34 in the last quarter; slope: β = −0.047, z = −2.26, p =.012). Baboons therefore recognized new exemplars of the trained category immediately. On negative trials, error was at chance in the first quarter (0.51, z = 0.91, p_FDR_ =.82). It fell significantly below chance from the second quarter onwards (0.43, z = −3.87, p_FDR_ <.001) and declined steeply over the batch (0.39 in the last quarter; slope: β = −0.163, z = −8.01, p <.001). Baboons therefore did not reject a new non-A category on first exposure; correct rejection was acquired through feedback within the batch.

Pooling across pairs could hide immediate generalization at late pairs, because negative generalization over the whole batch was absent at the earliest pairs (Table S2). We therefore repeated the analysis on pairs 5-8 only, where error on negative generalization trials was significantly below chance (Figure S1B). The pattern was unchanged. On negative trials, error in the first quarter was again at chance (0.50, z = 0.17, p_FDR_ =.61), then fell below chance from the second quarter onwards (0.41, z = −3.86, p_FDR_ <.001) and reached 0.34 in the last quarter (slope: β = −0.231, z = −8.35, p <.001). On positive trials, error was below chance in every quarter (first quarter: 0.38, z = −5.19, p_FDR_ <.001; last quarter: 0.31; slope: β = −0.104, z = −3.64, p <.001). Even after four or more pairs, therefore, a new non-A category was not rejected on first exposure. Within-batch slopes were numerically steeper at pairs 5-8 than in the pooled analysis for both stimulus types; this difference was not tested formally. That steepening fits savings, with feedback on a new pair being used more efficiently at late than at early pairs.

Negative stimuli thus benefited from previous training through savings rather than immediate generalization. Across pairs, the number of trials to criterion decreased (Figure 2A), and so did the error rate on negative generalization trials (Table S1). Each new non-A pair was therefore learned faster than the previous ones, although none was rejected on first exposure. This dissociation fits the familiarity account discussed in the main text. New positive exemplars belonged to the trained category and could be recognized at once. Each new negative exemplar belonged to a category never seen before, and what transferred across pairs was the speed of learning rather than an immediately applicable rule.

One limitation remains. At-chance error on negative trials combined with below-chance error on positive trials in the first quarter is also what a bias toward the “A” response would produce, and individual animals showed strong first-trial response biases. Average error across the two stimulus types in the first quarter (0.45 for pairs 2-8; 0.44 for pairs 5-8) indicates that discrimination was present but modest.

**Supplementary Figure S1.**
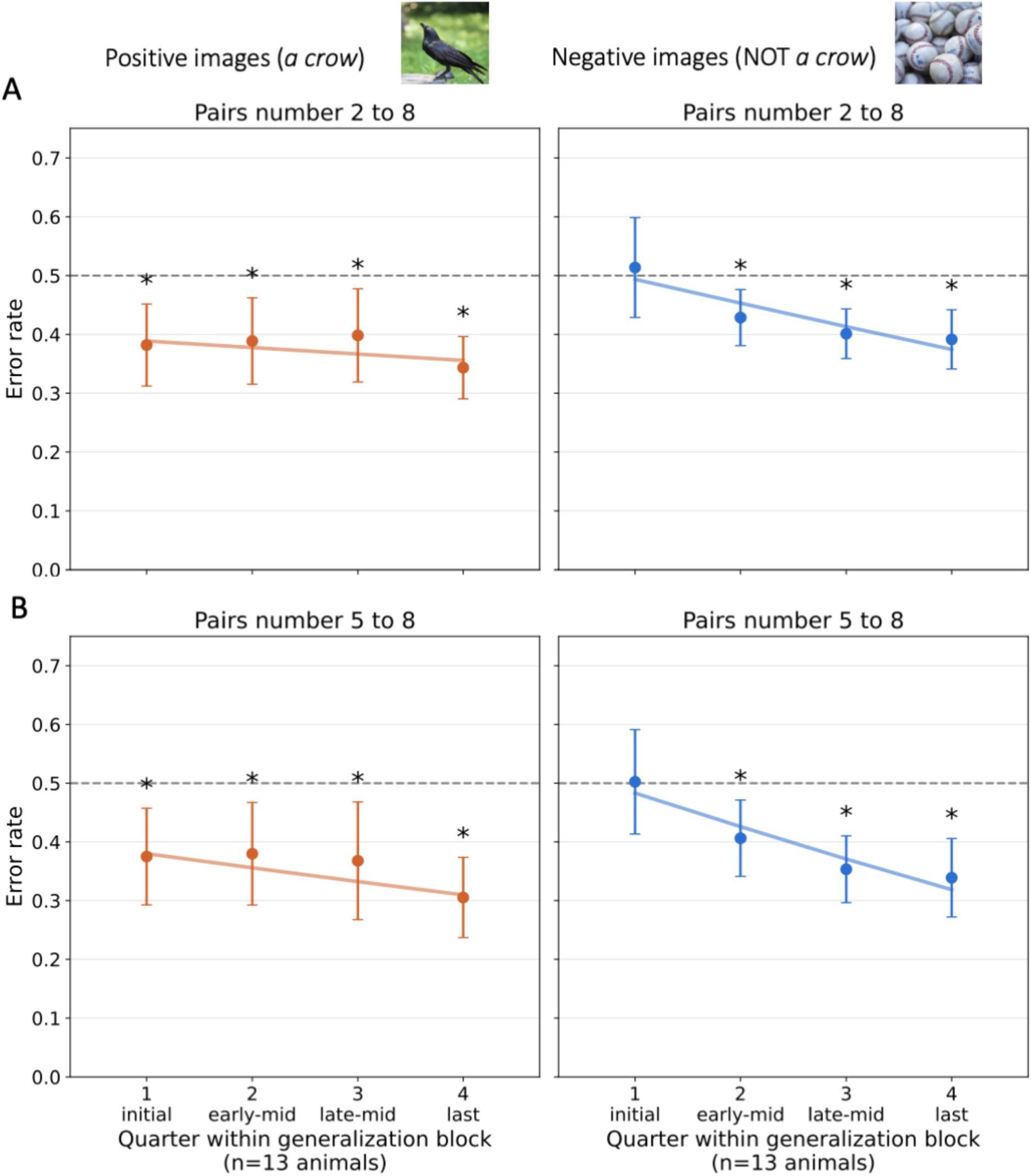
Immediate generalization for category A, but not for category not-A, in the naturalistic-image block. Error rate on generalization trials across four consecutive 8-trial quarters of the generalization batch (trials 1-8, 9-16, 17-24 and 25-32 of each stimulus type), for the 13 baboons that reached pair 8. Left panels show positive trials (orange) and right panels show negative trials (blue). (A) Data pooled across pairs 2-8. (B) Data restricted to pairs 5-8, where negative generalization over the whole batch was significantly below chance (Table S2). Circles are group means across animals ± 95% CI (1.96 × SEM). Lines are within-batch fits from a mixed-effects logistic regression of error on quarter, with a random intercept per animal. The dashed grey line marks chance (0.5). Asterisks mark quarters with error significantly below chance (one-sided GLMM tests, Benjamini−Hochberg FDR correction across the 16 quarters shown, p_FDR_<.05).

#### Both humans and baboons learn to categorize the color of geometrical stimuli

The color block served as a non-geometric perceptual control, but using the same line-pairs stimuli as the geometric categories. Unlike the block with naturalistic images, where categorization could draw on rich semantic and shape-based features for each category, the color block required participants to learn an abstract rule, i.e. deciding whether the two lines had the same color or not. This block allowed us to ask whether baboons’ ability to categorize natural images into A versus not-A would extend to an abstract rule. Note that color results are described first only for clarity, since the color and geometric blocks were actually intermixed in a random order, different for each participant.

#### Learning rat

The number of trials required to reach threshold rose from pair 1 (∼550 trials) to pairs 2 and 3 (∼950 trials) before declining sharply from pair 4 onwards to approximately 100 trials by pair 8 (exponential decay fit to pairs 2−8, R² = 0.877; Figure S2A). Human participants also learned the task efficiently, with trial counts near the imposed by our experimental mini-block procedure from pair 2 onwards (approximately 16−18 trials).

#### Error rate on generalization trials

For positive stimuli (Figure S2B), performance remained near chance across the first three pairs (error rates of approximately 0.49, 0.53 and 0.50), suggesting that the color rule was not accessible for generalization at these early stages, a markedly slower onset than for naturalistic images. Error rates then declined to approximately 0.37−0.40 at pairs 4 and 5, showed a transient increase at pair 6, and finally a sustained decline to approximately 0.30-0.35 at pairs 7 and 8. The fluctuation at pair 6 was also present among completers (Figures S3 and S4), whose composition does not change across pairs, and therefore cannot be explained by the reduction in sample size at that level alone. For negative stimuli, the trajectory was clearer and more pronounced: after fluctuating near chance across pairs 1-4, error rates dropped steeply from pair 5 onwards, reaching approximately 0.21 by pairs 7-8. Human participants showed near-floor generalization error rates from pair 2 onwards for both stimulus types (∼0.07 or below), with no meaningful asymmetry between positive and negative trials.

## Statistical analysis

For positive generalization trials, the error rate was significantly below chance at pairs 4 and 5, and again at pairs 7 and 8 (all FDR-adjusted p <.05; Table S2). For negative generalization trials, significant below-chance performance emerged earlier, at pairs 2 and 3, and again from pair 5 onwards. At pair 8, the error rate was estimated at 42.6% for positive stimuli (z = −22.04, p <.001) and at 27.9% for negative stimuli (z = −39.96, p <.001; Table S1). The learning slope was significantly negative for both positive (β = −0.051, z = −18.63, p <.001) and negative trials (β = −0.143, z = −29.83, p <.001), and steeper for the latter (trial-type × training-level interaction: β = −0.091, z = −23.36, p <.001). All these effects were preserved when the analysis was restricted to the six animals that completed the block (Tables S1−S2). Taken together, these results demonstrate that both baboons and humans can generalize a color-based rule to novel geometrical stimuli and, surprisingly, actually do better on negative trials than in positive ones. Note however that this finding merely means that they were better at judging two colors as different than as identical (see main text, discussion).

**Supplementary Figure S2.**
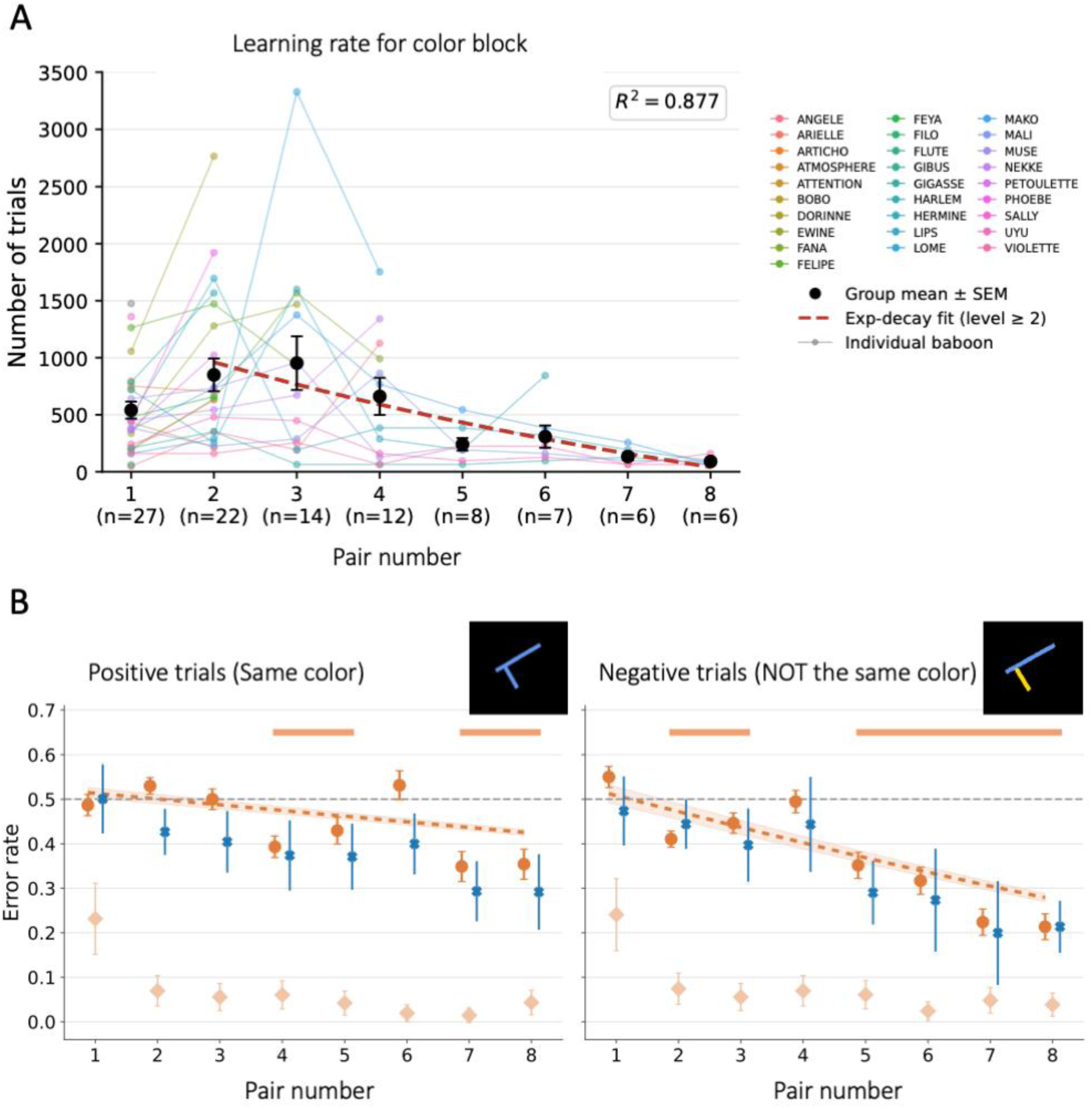
Baboons can categorize the color relation of the constituents of geometric stimuli. (A) Learning rate, as in Fig. 2A (exponential fit to levels 2-8, R² = 0.877). (B) Error rate on generalization trials for positive trials (left; two lines of the same color) and negative trials (right; two lines of different colors), as in Fig. 2B.

### Detailed generalization trajectories in the image and geometric blocks

#### Naturalistic images

In baboons, the error rate on positive generalization trials decreased steadily across pairs, reaching a plateau slightly below 40% from pair 4 onwards. The error rate on negative generalization trials also decreased, but plateaued later (around pair 6) and at a higher value (Figure 2B).

#### Perpendicularity

The generalization error rate on positive trials started near chance at pair 1 (∼0.55), then declined, reaching approximately 0.28−0.30 by pairs 6−7 before an upturn at pair 8 (∼0.41; Figure 3A). This upturn was also present among the six completers (Figure S5) and therefore reflects the specific stimulus pair rather than the reduction in sample size. For negative trials, after starting near chance at pair 1 (∼0.50), the error rate rose above chance at pairs 2−3 (∼0.55−0.57), before declining from pair 4 onwards to approximately 0.25 by pair 8.

#### Parallelism

The positive generalization error rate started near chance (∼0.55), declined irregularly across pairs, and reached approximately 0.40 at pair 8, with a spike back toward 0.56 at pair 7 (Figure 3B), observed in a small sample (n = 4). For negative trials, the decline was markedly steeper and earlier: starting near chance at pair 1 (∼0.48), the error rate dropped to approximately 0.38 by pair 3, fluctuated at pairs 4−6, then fell again to approximately 0.28−0.29 by pairs 7−8, ultimately reaching lower values than positive trials.

#### Length

The length block produced a distinct pattern, with a clear dissociation between positive and negative trials (Figure 3C). For positive trials, the generalization error rate declined steadily from pair 1 (∼0.57) to pairs 7−8 (∼0.27−0.34), one of the clearest and most regular learning trajectories observed across all blocks. For negative trials, by contrast, the error rate remained close to chance throughout, fluctuating between approximately 0.40 and 0.54 in the full sample, with no clear downward trend.

#### Model specification

The continuous-slope and categorical models agreed on all qualitative conclusions except for negative length trials, for which the continuous model estimated the error rate at chance at pair 8, whereas the categorical model detected small below-chance deviations at pairs 6 and 8 (Tables S1−S2). Among completers, both models detected below-chance performance on negative length trials at the last pairs.

### Robustness to attrition

#### Sample composition

The number of baboons contributing to each training level is reported in Table S3. Retention at pair 8 ranged from 50% (images) to 14% (parallelism and length), and the training history of each animal is given in Table S5. Because animals that fail to progress stop contributing data, any change in error rate across training levels in the full sample combines genuine learning with changes in the composition of the sample.

#### Pre-selection

To test whether completion was predicted by initial performance, we compared the error rate at pair 1, which all animals performed before any attrition, between animals that later completed the block and those that did not (Table S4). In a trial-level GLMM, completers made fewer errors than non-completers in all four blocks using geometrical stimuli (color: OR = 0.90; parallelism: OR = 0.91; angles: OR = 0.94; length: OR = 0.95; all FDR-adjusted p <.001), but not for naturalistic images (OR = 0.99, p =.78). At the level of per-animal error rates, the same trend was present (rank-biserial r = −.33 to −.43) but did not reach significance (Mann−Whitney tests, all FDR-adjusted p =.29), and pair-1 error rate correlated negatively with the maximum training level reached for color (p = −.55, p =.003), angles (p = −.47, p =.012) and parallelism (p = −.44, p =.018), but not for length (p = −.29, p =.13) or images (p = −.12, p =.56). The absolute difference was small (1−3 percentage points, with both groups close to chance), but indicates that attrition was partly performance-based.

#### Completers

We therefore refitted the continuous-slope and categorical models to the animals that reached pair 8 in each block (images: n = 13; color: n = 6; angles: n = 6; parallelism: n = 4; length: n = 4), for which the sample is identical at every pair (Tables S1−S2; Figures S3-S4). The error rate at pair 8 remained significantly below chance for positive stimuli in all five blocks, and for negative stimuli in the image, color, angles and parallelism blocks. The difference between positive and negative stimuli at pair 8 was preserved in all five blocks, in the same direction. The onset of positive generalization was preserved or occurred earlier (images: pair 2; color: pair 3; angles: pair 2; parallelism: pair 1; length: pair 2), as was the onset of negative generalization for images (pair 5) and parallelism (pair 3). Four effects differed between samples (Table S1). (i) For naturalistic images, the trial-type × training-level interaction was no longer significant (β = 0.000, z = 0.02, p =.98): in completers, positive and negative errors declined at the same rate (both β = −0.081), the negative category lagging behind by a constant offset: the cost of negation thus consists of a delayed onset and a constant offset rather than of a slower improvement. (ii) For parallelism, the slope on positive trials was no longer significant (β = +0.010, z = 1.65, p =.10), because completers were already below chance on positive stimuli at pairs 1 and 2. (iii) For angles, the trial-type × training-level interaction was no longer significant (β = −0.012, z = −1.71, p =.088). (iv) For length, the error rate on negative trials was significantly below chance at pair 8 (42.0%, z = −8.12, p <.001) and decreased significantly across pairs (β = −0.035, z = −3.44, p <.001), with below-chance performance sustained from pair 6 onwards. The positive−negative asymmetry for length remained large and significant (difference at pair 8: z = 21.09; interaction: z = 12.71; both p <.001). Because the geometric blocks were completed by only 4−6 animals, and because the GLMM standard errors primarily reflect trial-level variability, these estimates should be taken as characterizing the performance of these specific animals rather than of the baboon population.

**Supplementary Figure S3.**
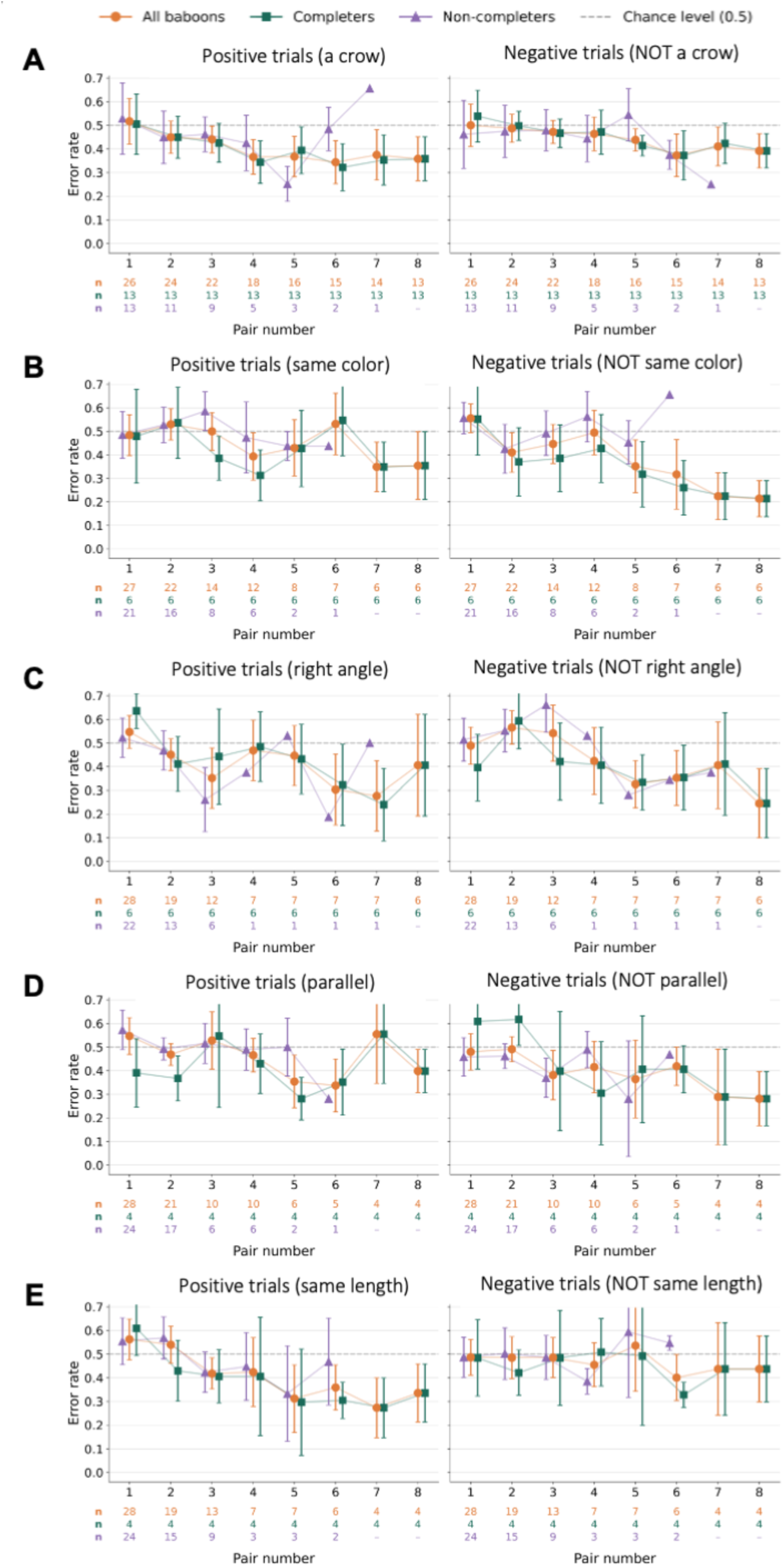
Generalization error in completers and non-completers. (A) Naturalistic images, (B) color, (C) perpendicularity, (D) parallelism, (E) length. Left subpanels show positive trials and right subpanels show negative trials. Orange circles are all baboons, green squares are baboons that completed the block, and purple triangles are baboons that did not (means across animals ± 95% CI, 1.96 × SEM). Rows below each subpanel give the number of baboons contributing at each training level.

**Supplementary Figure S4.**
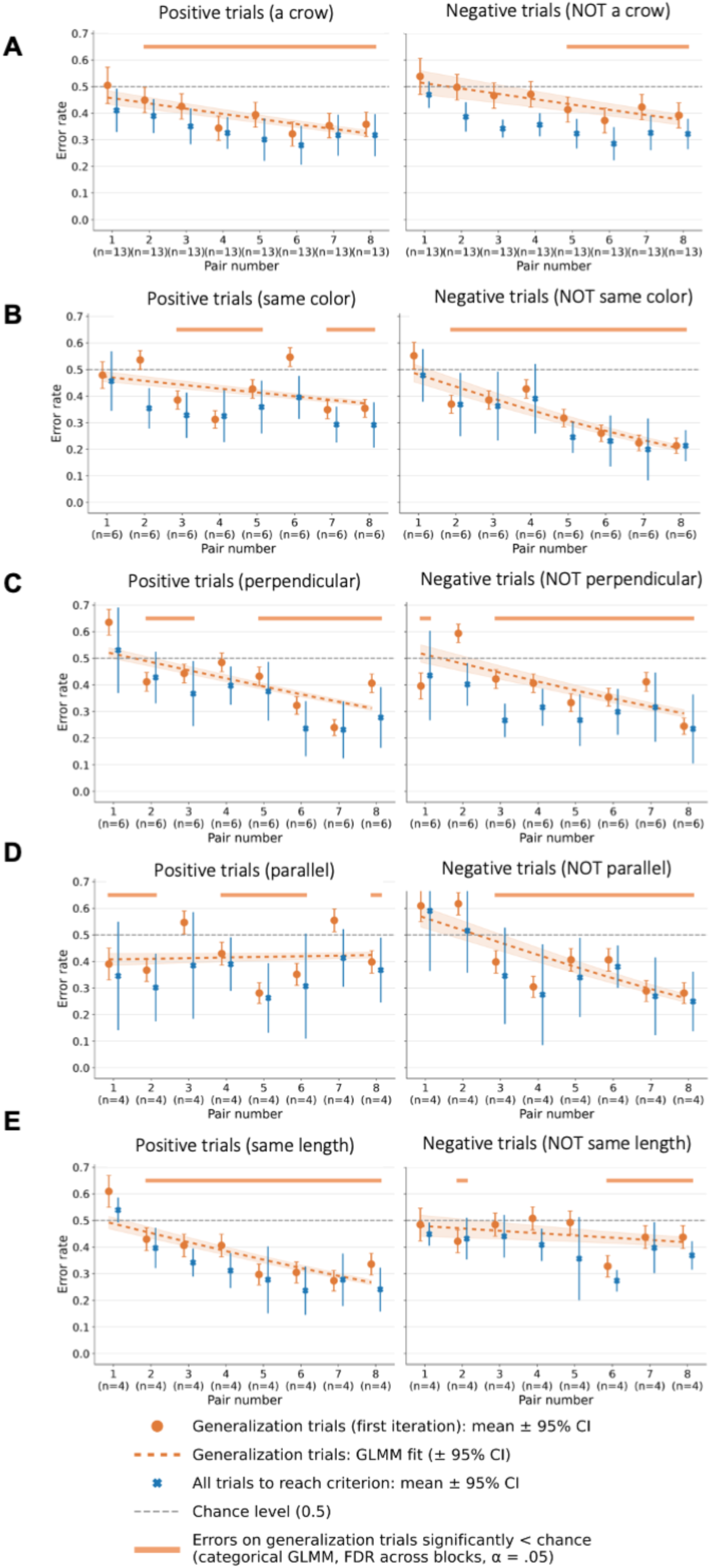
Generalization error restricted to baboons that completed each block. (A) Naturalistic images (n = 13), (B) color (n = 6), (C) perpendicularity (n = 6), (D) parallelism (n = 4), (E) length (n = 4). Left subpanels show positive trials and right subpanels show negative trials. Conventions are as in Fig. 2B, without human data. The GLMMs and FDR correction were refitted to the completers sample.

#### Training dose and intruder-task performance

Nine baboons performed the intruder task both before (2024) and after (2026) the geometry training, eight of which also took part in the original study. Their geometry training was heterogeneous (Table S6): they encountered between 5 and 19 geometric stimulus pairs (out of a maximum of 24) and completed between 0 and 2 of the three geometric blocks; across the whole cohort, no animal completed all three (Table S5). All nine animals improved between sessions (error-rate reduction: 0.01 to 0.21, Figure S5A). Neither measure of training dose correlated with the reduction in error rate, with the change in the symbolic coefficient, or with the change in the CorNet-IT coefficient (Figure S5B-C). For the number of pairs encountered, Spearman correlations were p = −.09 (improvement), p =.27 (Δβ symbolic) and p = −.06 (Δβ CorNet-IT), all p ≥.48; partial correlations controlling for baseline performance, total number of trials and age did not reach significance either (all p ≥.30), the largest being observed for the change in the symbolic coefficient (partial p =.42 to.46). For the number of geometric blocks completed, correlations were p =.15,.24 and −.01, respectively (all p ≥.53). Restricting the analysis to the eight animals common to all three sessions (including the initial Sablé-Meyer et al data) yielded the same pattern (all p ≥.15). At the individual level, the post-training symbolic coefficient did not reach significance in any animal after Holm correction; the largest was observed in one of the two animals that had completed two geometric blocks (β = 0.28, uncorrected p =.009, Holm-corrected p =.07). With nine animals, only correlations of |p| “.80 could be detected with 80% power; these analyses therefore rule out a strong dose−response relationship but cannot exclude a modest one.

**Supplementary Figure S5.**
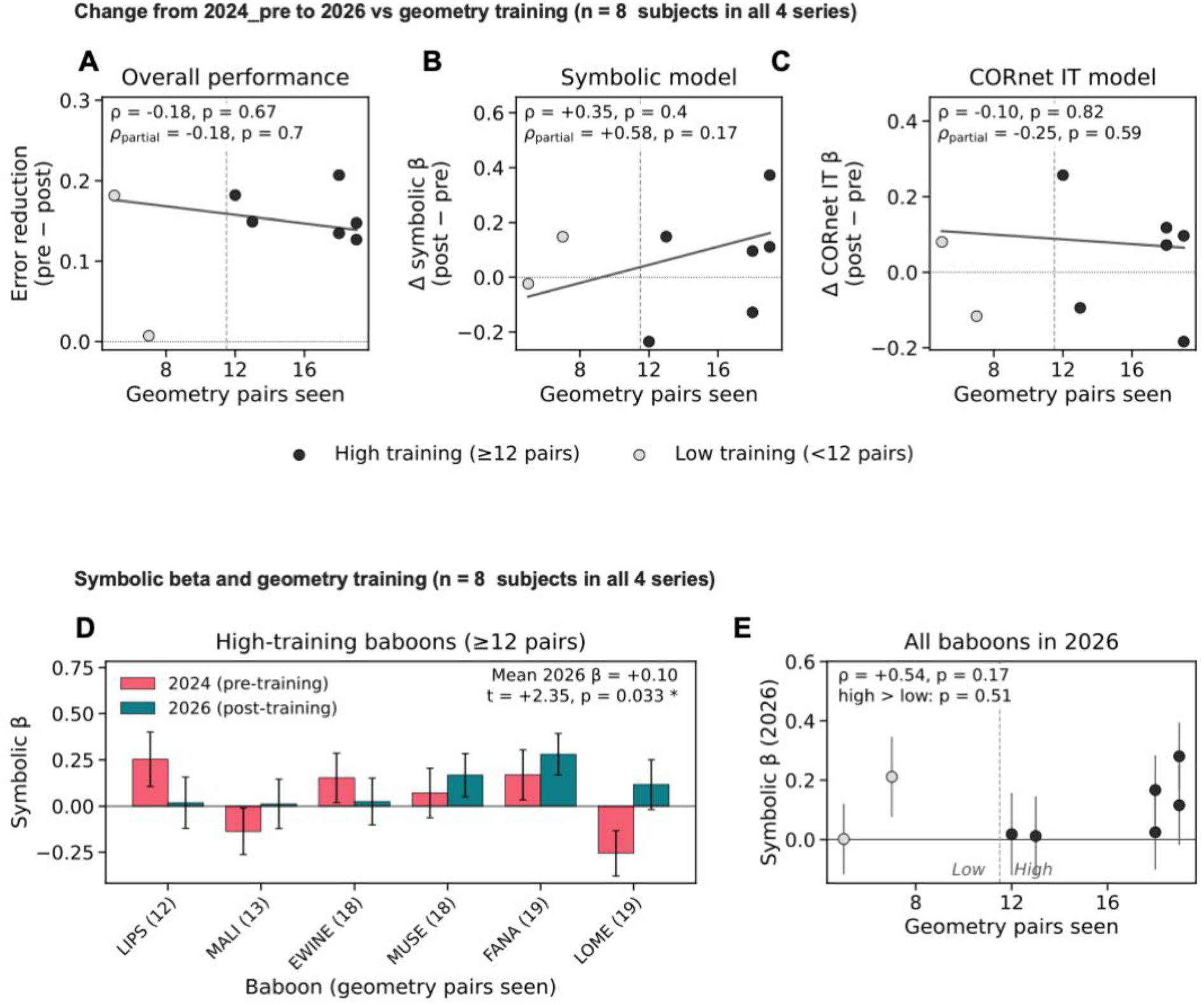
Amount of geometry training and change in intruder-task performance between 2024_pre and 2026, in the eight baboons also tested in 2019. • (A−C) Change as a function of the number of geometric stimulus pairs encountered: (A) error-rate reduction, Error(2024_pre) − Error(2026); (B) change in the symbolic coefficient; (C) change in the CorNet-IT coefficient. `o` Dots are individual baboons (grey, <12 pairs; black, ≥12 pairs). The solid line is the linear fit and the dotted line marks zero. `o` Spearman’s p and partial p controlling for baseline (two-sided); none is significant. The same analysis in all nine animals is reported in Supplementary Results. • (D) Symbolic coefficient in 2024_pre (pink) and 2026 (teal) for the six of these baboons with ≥12 pairs (number of pairs in brackets). Error bars are the SE of each coefficient. The 2026 group mean is +0.10 (one-sided one-sample t-test, t = 2.35, p = 0.033). •(E) 2026 symbolic coefficient of these 8 baboons as a function of pairs encountered. The dashed line marks the 12-pair threshold, and error bars are as in D. Spearman’s p = +0.54, p = 0.17 (two-sided). The comparison of ≥12 vs <12 pairs gives p = 0.51 (one-sided Welch t-test).

### Supplementary Methods

#### Housing and apparatus

The baboons lived in 3 groups. The first group of 20 animals lived in a 700-square-meter outdoor enclosure with access to 20-square-meter indoor housing. The second and third group, of 5 and 3 animals respectively, lived in a 30-square-meter (group 2) and 20-square-meter (group 3) outdoor enclosures with access to 10-square-meter (group 2) and 5-square-meter (group 3) indoor housings. They could engage, on a voluntary basis, with 14 automated learning devices located inside small covered cabins (10, 3 and 1 respectively for each group). Each device featured a 19-inch touch screen, a food dispenser, and a radio-frequency identification (RFID) reader that identified each animal. This identification allowed the experimenters to keep track of each animal’s progression through the experimental sessions and, at each new entrance into one of the cabins, to make sure that each baboon restarted from where it had left off. Overall, this setup provided a versatile and enriched environment, encouraging the baboons to interact with the learning devices at their own discretion.

### Stimuli and response symbols

#### Naturalistic stimuli

We selected 9 unique categories, each comprising 8 exemplars, from the THINGS initiative (43). For each baboon, one category was randomly assigned as the positive category (e.g., a crow), and one exemplar was drawn at random from each of the remaining 8 categories to serve as the negative stimulus of a specific pair (e.g., a baseball).

#### Geometrical stimuli

We used the drawvsg Python library to generate a set of baboon-specific geometrical stimuli. Each stimulus consisted of two lines that were either secant or non-secant, and could vary along three independent axes: (i) parallelism and angles (i.e., being parallel or not parallel; forming a right angle or not), (ii) the relative length of the two lines (i.e., being equal or unequal), and (iii) the color of the two lines (i.e., being the same or different). Stimuli were organized into pairs such that the two members of a pair differed on exactly one axis − the axis corresponding to the concept under test − while being maximally dissimilar from the pairs used at other training levels. A given stimulus (for instance, parallel green lines of the same length) could thus fulfill several properties (in this example: parallelism, same color and same length). Non-parallel lines subtended angles of ±30°, ±40°, ±50° or ±60°; non-right-angled lines subtended the same set of angular deviations from 90°. The color palette comprised nine distinct colors: white, dark green, orange-red, gold, crimson, navy blue, lime green, cyan and cornflower blue. To prevent baboons from exploiting low-level perceptual regularities, two task-irrelevant parameters were also varied for each stimulus: its global orientation (randomly drawn from 30° to 360°, excluding 90°, 180° and 270°) and the absolute length of the lines (varied by up to ±30% from the reference version).

#### Response symbols

The response options consisted of abstract symbols inspired by the lexigrams used with the bonobo Kanzi (51). Symbols were deliberately designed to exclude straight lines and sharp angles, to avoid any visual confound with the geometrical properties under test. A total of 10 symbols were created and grouped into 5 maximally dissimilar pairs using a CLIP-based perceptual distance (52). For each baboon and each block, one pair of symbols was randomly assigned, the two symbols serving, respectively, as the positive and negative response options throughout the block.

#### Structure of the task

Each baboon completed the five blocks in a randomized order. Within each block, the concept was introduced progressively across up to eight training levels. At each new training level, a novel stimulus pair was introduced, and the baboon was tested exclusively on that pair until reaching the success criterion of 70% correct responses on both positive and negative stimuli. The first training level comprised batches of 32 trials, and all subsequent levels batches of 64 trials; this asymmetry was meant to let baboons fully engaged in the task move on to the second pair as rapidly as possible. A single pair of response symbols was used throughout each block, and the position of the symbols on the screen (left or right) was randomized across trials to prevent response biases. Expecting that humans would rapidly reach near-ceiling performance, the batches were reduced to 8 trials for the first training level and 16 trials for subsequent levels.

Experiments were programmed using the OpenSesame software (53) and the Open Monkey Mind extension.

#### Analysis of learning rate

For each task block, learning rate was quantified as the number of trials required to reach the success criterion for each pair of stimuli, derived from the number of completed batches (32 trials per batch for pair 1, 64 trials for pairs 2 to 8). For each participant and pair, trial counts were averaged across experimental sessions. Group-level performance for each pair was then summarized as the mean across subjects ± the standard error of the mean (SEM), and individual trajectories were overlaid to visualize inter-subject variability.

To characterize the shape of the group-level learning curve, we compared four candidate parametric families − linear, logarithmic, power, and three-parameter exponential decay − fitted to the subject-averaged means by nonlinear least squares (Levenberg−Marquardt algorithm, scipy.optimize.curve_fit). Goodness of fit was quantified as the coefficient of determination (R²) computed on the group means. Across blocks, the three-parameter exponential decay model y = a·exp(−b·x) + c, in which a captures the initial offset above asymptote, b the learning rate, and c the asymptotic number of trials, consistently provided the best or near-best fit and was therefore adopted as the common functional form for all blocks. For numerical stability, initial parameter estimates were derived from the data (co = final observed value, ao = initial minus final value, bo = 1/range of training levels) and bounded to b ≥ 0 and c ≥ 0 to enforce a non-increasing curve with a non-negative asymptote. All analyses were performed in Python using NumPy, pandas, SciPy and scikit-learn; figures were produced with Matplotlib and Seaborn.

### Statistical analysis of generalization

#### Data

We analyzed baboon errors (1 = error, 0 = correct) on generalization trials, that is, the first batch of each training level (32 trials at pair 1, 64 at later pairs), in which the new stimulus pair was presented for the first time. Each block was analyzed separately.

#### Model fitting

All models were binomial generalized linear mixed models (logit link) with a random intercept per animal, fitted by variational Bayes (BinomialBayesMixedGLM, statsmodels; default priors: normal with SD = 2 on fixed effects, SD = 1 on the log standard deviation of the random intercept). For each estimate we report the posterior mean (B) and standard deviation (SD), with z = B/SD and an approximate p-value from the normal distribution. For quantities that combine several coefficients, the SD was obtained by summing their variances, as the variational posterior treats coefficients as independent. Predicted error rates are for an average animal (random intercept = 0).

#### Continuous model

The main model was errors ∼ training level × trial type + (1 | animal). Training level was centered on pair 8 (level − 8), so that the intercept is the log-odds of error at pair 8, and positive trials were the reference. This single model answered three questions per block, with two-sided tests (Table S1):

1. Does error at pair 8 differ from chance (log-odds = 0, i.e. 50% error)? Intercept for positive trials; intercept + trial-type effect for negative trials.
2. Does error decrease across pairs? Level slope for positive trials; slope + interaction for negative trials.
3. Do positive and negative trials differ? Trial-type effect (difference at pair 8) and interaction (difference in slope).

The fitted curves and 95% credible bands in Figs. 2B, 3 and S4 are this model’s predictions (posterior mean ± 1.96 SD on the log-odds scale, converted to error rates).

#### Categorical model

To find the pairs at which error was below chance without assuming a linear trend, we refitted each block with training level as a categorical factor (pair 8 as reference) in place of the slope. The log-odds of error at each pair and trial type was obtained by adding the relevant coefficients and tested with a one-sided test (error < 50%). The resulting 80 tests (5 blocks × 8 pairs × 2 trial types) were corrected together with the Benjamini−Hochberg procedure (a =.05). These corrected tests define the generalization onsets in the main text, the below-chance markers in the figures, and Table S2.

### Attrition analyses

#### Completers

For each block, completers were defined as the animals that reached the last training level (pair 8). The continuous-slope and categorical GLMMs described above were refitted to the generalization trials of these animals only, with the same fixed and random effects. For the categorical model, the Benjamini−Hochberg procedure was applied across the 80 tests of the completers sample, as for the full sample. A conclusion was considered to differ between samples when an effect crossed the a =.05 threshold in either direction, or when a significant effect changed sign.

#### Pre-selection

For each block, we compared the pair-1 error rate (all trials) of completers and non-completers using (i) Mann−Whitney tests on per-animal error rates, with the rank-biserial correlation as effect size, (ii) Spearman correlations between per-animal pair-1 error rate and the maximum training level reached, and (iii) a trial-level binomial GLMM with completion status as a fixed effect and a random intercept per animal. p-values from (i) and (iii) were corrected across the five blocks using the Benjamini−Hochberg procedure.

#### Quadrilateral intruder task

The task used a set of 11 fixed quadrilateral reference shapes, matched for overall size (equalized mean pairwise vertex distance) and, for 9 of the 11 shapes, for bottom-edge length. For each reference shape, four deviant shapes were generated by repositioning the bottom-right vertex at a fixed distance from its correct location (30% of the mean pairwise vertex distance): two deviants altered the length of the bottom edge, and two preserved its length but changed its orientation. To prevent participants from exploiting screen-relative parallelism cues or rotational symmetry, shapes were presented at one of six randomly assigned orientations (±5°, ±15°, ±25° from canonical) and one of six randomly assigned size scaling factors (0.875 to 1.125 applied to edge lengths), each value being used exactly once across the six shapes of a trial. Each block comprised 88 trials (one per reference shape × deviant type × outlier position combination).

The three iterations of the task differed in length: baboons completed up to 99 blocks in the initial study, up to 60 blocks in the pre-training replication, and up to 30 blocks in the post-training test. Because performance improves with practice within a session, all cross-session comparisons were restricted to the first 30 blocks per animal (≈2,640 trials; all trials for animals with fewer), so that sessions reflected a comparable amount of exposure to the task. The version of the initial-study data used for this matched comparison (“2019_matched”) consisted of the first 30 blocks of each animal’s session, whereas the previously reported “2019” data corresponded to the most-practiced portion of that longer session.

#### Performance analysis

Performance was quantified, for each animal, as the proportion of erroneous responses (error rate) per reference shape, where an error was any selection of a non-deviant shape. Error rates were computed per animal and per session and averaged across the eight animals that completed all three sessions for the group-level curves (Figure 4A). Group-level error rates are reported as the mean across subjects ± SEM. Within-session practice effects were assessed by comparing the first and last blocks of the initial-study session; cross-session differences in overall error rate were evaluated on the trial-matched data.

#### Computational modelling

To assess which representations best accounted for the pattern of errors across conditions, we followed the modelling approach of Sablé-Meyer et al (30). and compared a symbolic model, which encodes each shape by a compressed geometric description as a sum of its basic geometric properties, with a non-symbolic model based on the IT-layer activations of the CorNet-S convolutional neural network (45). For each model, a single predicted difficulty value was derived for each of the 44 conditions (11 reference shapes × 4 deviant shapes), z-scored across conditions, and sign-flipped so that higher values corresponded to conditions predicted to be harder. For each animal and session, we then fitted an ordinary-least-squares regression of the z-scored error rate across conditions on the two predictors entered jointly (error ∼ symbolic + CorNet-IT, no intercept), yielding one coefficient (β) per model. The per-animal β values were tested against zero at the session level using one-sided one-sample t-tests (H1: β > 0), and were compared between sessions in the animals that performed both sessions (one-sided paired tests: symbolic, H1 increase; CorNet-IT, H1 decrease), the symbolic and CorNet-IT changes being contrasted by a difference-of-difference test. These analyses were performed on all animals with available data in each session (n = 10−20 per session; n = 9−13 per pairwise comparison). For the individual post-training symbolic coefficients, p-values were corrected across animals using the Holm procedure.

#### Training-dose analysis

The analysis included all animals that performed both the pre-training (2024) and the post-training (2026) intruder sessions (n = 9), and was repeated on the subset that also performed the initial session (n = 8). Training dose was quantified as the total number of stimulus pairs encountered across the angles, parallelism and length blocks (maximum 24) and, alternatively, as the number of these blocks completed. Outcomes were computed on the trial-matched data: the improvement in overall error rate (pre − post) and the change (post − pre) in the per-animal symbolic and CorNet-IT coefficients. Associations were assessed with Spearman and Pearson correlations and with partial Spearman correlations controlling successively for the pre-training value of the outcome, the total number of trials performed across the categorization blocks, and age.

#### Within-batch analysis of generalization trials (image block)

We tested whether below-chance error on generalization trials was present from the first presentations of a new pair, or emerged within the batch through feedback. For each baboon and training level (pairs 2-8; pair 1 was excluded because it corresponds to task acquisition), we took the 32 positive and 32 negative generalization trials in chronological order. We then split each sequence into four consecutive quarters of 8 trials (trials 1-8, 9-16, 17-24, 25-32). The analysis was restricted to the 13 baboons that reached pair 8, so that every quarter and every pair included the same animals. Plotted error rates were averaged within each animal and then across animals.

Two tests were carried out with binomial GLMMs with a random intercept per animal, fitted separately for positive and negative trials. First, to test whether each quarter was below chance, quarter was entered as a categorical factor, and the model-estimated log-odds of error in each quarter was tested against zero (error rate=.5) with a one-sided test. Benjamini−Hochberg FDR correction was applied across the 16 quarter estimates of the figure (2 pair ranges × 2 trial types × 4 quarters). Second, to test whether error declined within the batch, quarter index (1-4) was entered as a continuous predictor, and its slope was tested with a one-sided test (H1: slope < 0), with Benjamini−Hochberg correction across the two trial types.

Because negative generalization over the whole batch emerged only from pair 5 (Table S2), pooling across pairs 2-8 could hide immediate generalization at later pairs. We therefore repeated the same analysis, with the same animals, tests and models, on pairs 5-8 only, for both stimulus types.

---

**Table S1.** Generalization error on first-iteration trials as a function of training level, in all animals and in completers. Errors were modeled with binomial GLMMs (logit link, variational Bayes) including training level (continuous, centered on level 8), trial type (positive, negative) and their interaction, with a random intercept per animal. Models were fitted separately for each block, first on all animals that performed the block and then on completers only (animals that reached level 8 in that block). The *n* for each sample is given in each block header. “vs. chance” rows test the predicted log-odds of error at level 8 against 0 (error rate =.50), and *P*(error) is the corresponding predicted error rate. “Negative − positive” rows test the trial-type difference at level 8 and the trial type × training level interaction. Slopes are per training level, and negative values indicate decreasing error. *B*, posterior mean (log-odds of error); *SD*, posterior standard deviation; *z* = *B*/*SD*; *p*, two-sided normal approximation, uncorrected. ^a^ Conclusion at a =.05 differs between the two samples.

|  | All animals |  |  |  |  | Completers |  |  |  |  |
|---|---|---|---|---|---|---|---|---|---|---|
| Effect | *P*(error) | *B* | *SD* | *z* | *p* | *P*(error) | *B* | *SD* | *z* | *p* |
| Naturalistic images (all animals: *n* = 26; completers: *n* = 13) |  |  |  |  |  |  |  |  |  |  |
| Level 8: positive vs. chance | .33 | −0.710 | 0.022 | −32.51 | < .001 | .32 | −0.731 | 0.026 | −28.35 | < .001 |
| Level 8: negative vs. chance | .38 | −0.478 | 0.038 | −12.71 | < .001 | .38 | −0.504 | 0.044 | −11.35 | < .001 |
| Level 8: negative − positive | — | 0.231 | 0.031 | 7.55 | < .001 | — | 0.227 | 0.036 | 6.28 | < .001 |
| Slope: positive | — | −0.087 | 0.005 | −17.49 | < .001 | — | −0.081 | 0.006 | −12.49 | < .001 |
| Slope: negative | — | −0.071 | 0.009 | −8.29 | < .001 | — | −0.081 | 0.011 | −7.21 | < .001 |
| Slope: negative − positive | — | 0.016 | 0.007 | 2.24 | .025 | — | 0.000 | 0.009 | 0.02 | .982<sup>a</sup> |
| Color (all animals: *n* = 27; completers: *n* = 6) |  |  |  |  |  |  |  |  |  |  |
| Level 8: positive vs. chance | .43 | −0.300 | 0.014 | −22.04 | < .001 | .37 | −0.518 | 0.020 | −26.54 | < .001 |
| Level 8: negative vs. chance | .28 | −0.949 | 0.024 | −39.96 | < .001 | .21 | −1.346 | 0.035 | −38.93 | < .001 |
| Level 8: negative − positive | — | −0.650 | 0.019 | −33.35 | < .001 | — | −0.828 | 0.029 | −29.01 | < .001 |
| Slope: positive | — | −0.051 | 0.003 | −18.63 | < .001 | — | −0.059 | 0.005 | −12.28 | < .001 |
| Slope: negative | — | −0.143 | 0.005 | −29.83 | < .001 | — | −0.185 | 0.008 | −22.00 | < .001 |
| Slope: negative − positive | — | −0.091 | 0.004 | −23.36 | < .001 | — | −0.126 | 0.007 | −18.26 | < .001 |
| Right angles (all animals: *n* = 28; completers: *n* = 6) |  |  |  |  |  |  |  |  |  |  |
| Level 8: positive vs. chance | .32 | −0.733 | 0.014 | −50.69 | < .001 | .31 | −0.788 | 0.019 | −40.97 | < .001 |
| Level 8: negative vs. chance | .31 | −0.806 | 0.025 | −32.23 | < .001 | .29 | −0.882 | 0.033 | −26.39 | < .001 |
| Level 8: negative − positive | — | −0.074 | 0.020 | −3.62 | < .001 | — | −0.093 | 0.027 | −3.42 | < .001 |
| Slope: positive | — | −0.098 | 0.003 | −33.47 | < .001 | — | −0.124 | 0.005 | −25.95 | < .001 |
| Slope: negative | — | −0.152 | 0.005 | −29.83 | < .001 | — | −0.136 | 0.008 | −16.36 | < .001 |
| Slope: negative − positive | — | −0.053 | 0.004 | −12.87 | < .001 | — | −0.012 | 0.007 | −1.71 | .088<sup>a</sup> |
| Parallelism (all animals: *n* = 28; completers: *n* = 4) |  |  |  |  |  |  |  |  |  |  |
| Level 8: positive vs. chance | .41 | −0.382 | 0.015 | −25.83 | < .001 | .42 | −0.306 | 0.023 | −13.05 | < .001 |
| Level 8: negative vs. chance | .31 | −0.805 | 0.026 | −31.25 | < .001 | .26 | −1.029 | 0.041 | −25.14 | < .001 |
| Level 8: negative − positive | — | −0.423 | 0.021 | −20.05 | < .001 | — | −0.723 | 0.034 | −21.54 | < .001 |
| Slope: positive | — | −0.064 | 0.003 | −22.01 | < .001 | — | 0.010 | 0.006 | 1.65 | .099<sup>a</sup> |
| Slope: negative | — | −0.113 | 0.005 | −22.50 | < .001 | — | −0.187 | 0.010 | −18.37 | < .001 |
| Slope: negative − positive | — | −0.049 | 0.004 | −12.00 | < .001 | — | −0.197 | 0.008 | −23.69 | < .001 |
| Length (all animals: *n* = 28; completers: *n* = 4) |  |  |  |  |  |  |  |  |  |  |
| Level 8: positive vs. chance | .31 | −0.810 | 0.015 | −54.91 | < .001 | .27 | −1.009 | 0.023 | −43.14 | < .001 |
| Level 8: negative vs. chance | .50 | 0.006 | 0.025 | 0.23 | .815 | .42 | −0.325 | 0.040 | −8.12 | < .001<sup>a</sup> |
| Level 8: negative − positive | — | 0.816 | 0.021 | 39.50 | < .001 | — | 0.684 | 0.032 | 21.09 | < .001 |
| Slope: positive | — | −0.144 | 0.003 | −49.85 | < .001 | — | −0.140 | 0.006 | −23.83 | < .001 |
| Slope: negative | — | 0.009 | 0.005 | 1.81 | .071 | — | −0.035 | 0.010 | −3.44 | < .001<sup>a</sup> |
| Slope: negative − positive | — | 0.153 | 0.004 | 37.55 | < .001 | — | 0.105 | 0.008 | 12.71 | < .001 |

*Note.* Binomial GLMMs (logit link, variational Bayes) of first-iteration errors: errors ~ training level (centered at 8) × trial type + (1 \| animal), fitted separately for each block. Completers are the animals that reached training level 8 in that block. *B* = posterior mean (log-odds of error); *SD* = posterior standard deviation; *z* = *B*/*SD*; *p* = two-sided, normal approximation, uncorrected. “vs. chance” rows test the model-predicted log-odds at level 8 against 0 (error rate = .50); *P*(error) is the corresponding predicted error rate. “Negative − positive” rows test the trial-type main effect at level 8 and the trial type × training level interaction. Slopes are per training level; negative slopes indicate decreasing error.
<sup>a</sup> Inferential conclusion at α = .05 differs between the two samples.

**Table S2.**
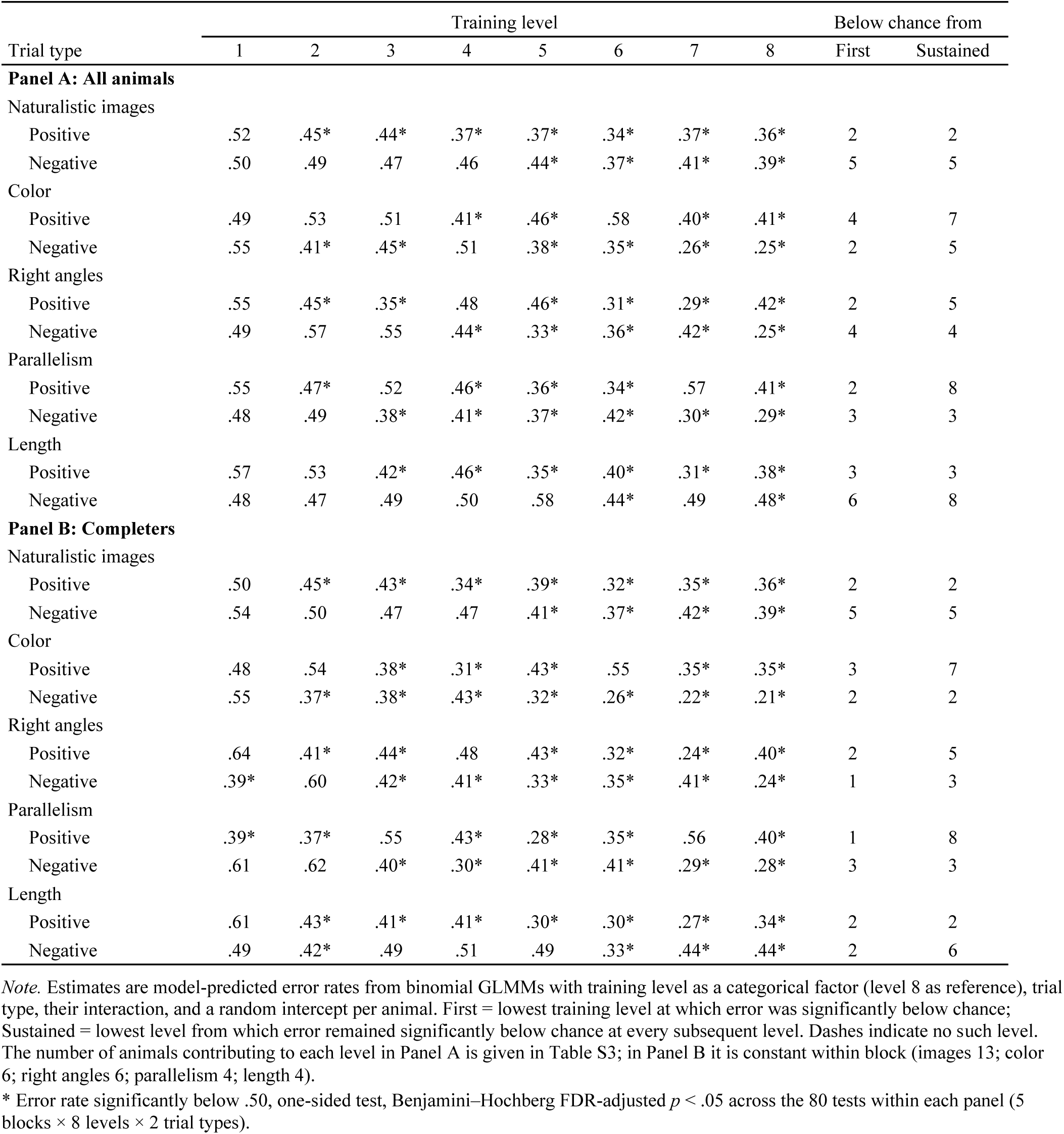
Model-estimated error rate on generalization trials at each training level. Predicted error rates come from binomial GLMMs treating training level as a categorical factor (reference: level 8), with trial type, their interaction and a random intercept per animal. They were fitted separately for each block in (*A*) all animals and (*B*) completers. *Error rate significantly below chance (.50): one-sided tests, Benjamini– Hochberg FDR-adjusted *p* <.05 across the 80 tests within each panel (5 blocks × 8 levels × 2 trial types). “First” is the lowest level at which error was significantly below chance. “Sustained” is the lowest level from which error remained significantly below chance at every subsequent level. The number of animals contributing to each level is given in Table S3 for panel *A*. In panel *B* it is constant within each block: naturalistic images, 13; color, 6; right angles, 6; parallelism, 4; length, 4.

|  | Training level |  |  |  |  |  |  |  | Below chance from |  |
|---|---|---|---|---|---|---|---|---|---|---|
| Trial type | 1 | 2 | 3 | 4 | 5 | 6 | 7 | 8 | First | Sustained |
| **Panel A: All animals** |  |  |  |  |  |  |  |  |  |  |
| Naturalistic images |  |  |  |  |  |  |  |  |  |  |
| Positive | .52 | .45* | .44* | .37* | .37* | .34* | .37* | .36* | 2 | 2 |
| Negative | .50 | .49 | .47 | .46 | .44* | .37* | .41* | .39* | 5 | 5 |
| Color |  |  |  |  |  |  |  |  |  |  |
| Positive | .49 | .53 | .51 | .41* | .46* | .58 | .40* | .41* | 4 | 7 |
| Negative | .55 | .41* | .45* | .51 | .38* | .35* | .26* | .25* | 2 | 5 |
| Right angles |  |  |  |  |  |  |  |  |  |  |
| Positive | .55 | .45* | .35* | .48 | .46* | .31* | .29* | .42* | 2 | 5 |
| Negative | .49 | .57 | .55 | .44* | .33* | .36* | .42* | .25* | 4 | 4 |
| Parallelism |  |  |  |  |  |  |  |  |  |  |
| Positive | .55 | .47* | .52 | .46* | .36* | .34* | .57 | .41* | 2 | 8 |
| Negative | .48 | .49 | .38* | .41* | .37* | .42* | .30* | .29* | 3 | 3 |
| Length |  |  |  |  |  |  |  |  |  |  |
| Positive | .57 | .53 | .42* | .46* | .35* | .40* | .31* | .38* | 3 | 3 |
| Negative | .48 | .47 | .49 | .50 | .58 | .44* | .49 | .48* | 6 | 8 |
| **Panel B: Completers** |  |  |  |  |  |  |  |  |  |  |
| Naturalistic images |  |  |  |  |  |  |  |  |  |  |
| Positive | .50 | .45* | .43* | .34* | .39* | .32* | .35* | .36* | 2 | 2 |
| Negative | .54 | .50 | .47 | .47 | .41* | .37* | .42* | .39* | 5 | 5 |
| Color |  |  |  |  |  |  |  |  |  |  |
| Positive | .48 | .54 | .38* | .31* | .43* | .55 | .35* | .35* | 3 | 7 |
| Negative | .55 | .37* | .38* | .43* | .32* | .26* | .22* | .21* | 2 | 2 |
| Right angles |  |  |  |  |  |  |  |  |  |  |
| Positive | .64 | .41* | .44* | .48 | .43* | .32* | .24* | .40* | 2 | 5 |
| Negative | .39* | .60 | .42* | .41* | .33* | .35* | .41* | .24* | 1 | 3 |
| Parallelism |  |  |  |  |  |  |  |  |  |  |
| Positive | .39* | .37* | .55 | .43* | .28* | .35* | .56 | .40* | 1 | 8 |
| Negative | .61 | .62 | .40* | .30* | .41* | .41* | .29* | .28* | 3 | 3 |
| Length |  |  |  |  |  |  |  |  |  |  |
| Positive | .61 | .43* | .41* | .41* | .30* | .30* | .27* | .34* | 2 | 2 |
| Negative | .49 | .42* | .49 | .51 | .49 | .33* | .44* | .44* | 2 | 6 |

*Note.* Estimates are model-predicted error rates from binomial GLMMs with training level as a categorical factor (level 8 as reference), trial type, their interaction, and a random intercept per animal. First = lowest training level at which error was significantly below chance; Sustained = lowest level from which error remained significantly below chance at every subsequent level. Dashes indicate no such level. The number of animals contributing to each level in Panel A is given in Table S3; in Panel B it is constant within block (images 13; color 6; right angles 6; parallelism 4; length 4).
\* Error rate significantly below .50, one-sided test, Benjamini–Hochberg FDR-adjusted *p* < .05 across the 80 tests within each panel (5 blocks × 8 levels × 2 trial types).

**Table S3.** Number of baboons contributing data at each training level. An animal contributes to a level once it has reached it. Animals reaching level 8 form the completers sample of Tables S1 and S2. The last column gives the percentage of animals starting the block that reached level 8. One animal did not perform the naturalistic-image block, and another performed neither the image nor the color block (Table S5), which lowers the starting *n* for those blocks.

|  | Training level |  |  |  |  |  |  |  |  |
|---|---|---|---|---|---|---|---|---|---|
| Block | 1 | 2 | 3 | 4 | 5 | 6 | 7 | 8 | % retained at level 8 |
| Naturalistic images | 26 | 24 | 22 | 18 | 16 | 15 | 14 | 13 | 50 |
| Color | 27 | 22 | 14 | 12 | 8 | 7 | 6 | 6 | 22 |
| Right angles | 28 | 19 | 12 | 7 | 7 | 7 | 7 | 6 | 21 |
| Parallelism | 28 | 21 | 10 | 10 | 6 | 5 | 4 | 4 | 14 |
| Length | 28 | 19 | 13 | 7 | 7 | 6 | 4 | 4 | 14 |

*Note.* An animal contributes to a level once it has reached it. Animals reaching level 8 constitute the completers sample (Tables S1 and S2). One animal did not perform the image block and another performed neither the image nor the color block (Table S5).

**Table S4.** Level-1 error rates of animals that later did or did not complete each block. All animals performed level 1 before any attrition, so these comparisons test whether completing a block was predicted by initial performance. C, completers (reached level 8); NC, non-completers. Mann–Whitney tests compare per-animal level-1 error rates. *r*∼rb∼ is the rank-biserial correlation, where negative values mean lower error in completers. p is the Spearman correlation between level-1 error and the maximum level reached in the block (uncorrected). The GLMM is a trial-level binomial model with completion status as predictor and a random intercept per animal. *B*, posterior mean (log-odds); *OR*, odds ratio (< 1: lower odds of error in completers) with 95% credible interval (CrI). *p*∼FDR∼, Benjamini–Hochberg adjustment across the five blocks.

|  | Sample |  | Mean level-1 error |  | Mann–Whitney |  |  | Error vs. max level |  | GLMM: completer vs. non-completer |  |  |  |
|---|---|---|---|---|---|---|---|---|---|---|---|---|---|
| Block | *n*<sub>C</sub> | *n*<sub>NC</sub> | C | NC | *U* | *r*<sub>rb</sub> | *p*<sub>FDR</sub> | ρ | *p* | *B* | *z* | *OR* [95% CrI] | *p*<sub>FDR</sub> |
| Naturalistic images | 13 | 13 | .44 | .45 | 88.0 | .04 | .878 | −.12 | .564 | −0.008 | −0.28 | 0.99 [0.94, 1.05] | .782 |
| Color | 6 | 21 | .47 | .49 | 36.0 | −.43 | .294 | −.55 | .003 | −0.104 | −6.14 | 0.90 [0.87, 0.93] | < .001 |
| Right angles | 6 | 22 | .48 | .49 | 44.0 | −.33 | .294 | −.47 | .012 | −0.061 | −5.16 | 0.94 [0.92, 0.96] | < .001 |
| Parallelism | 4 | 24 | .47 | .49 | 28.0 | −.42 | .294 | −.44 | .018 | −0.095 | −6.30 | 0.91 [0.88, 0.94] | < .001 |
| Length | 4 | 24 | .49 | .51 | 29.0 | −.40 | .294 | −.29 | .131 | −0.049 | −3.40 | 0.95 [0.93, 0.98] | < .001 |

*Note.* Level 1 was performed by all animals before any attrition, so these comparisons test whether completion was predicted by initial performance. C = completers (reached level 8); NC = non-completers. Mann–Whitney tests compare per-animal error rates; *r*<sub>rb</sub> = rank-biserial correlation (negative values: lower error in completers). ρ = Spearman correlation between level-1 error and the maximum level reached (uncorrected). The GLMM is a trial-level binomial model with a random intercept per animal; *OR* < 1 indicates lower odds of error in completers; CrI = credible interval. *p*<sub>FDR</sub> = Benjamini–Hochberg adjustment across the five blocks.

**Table S5.**
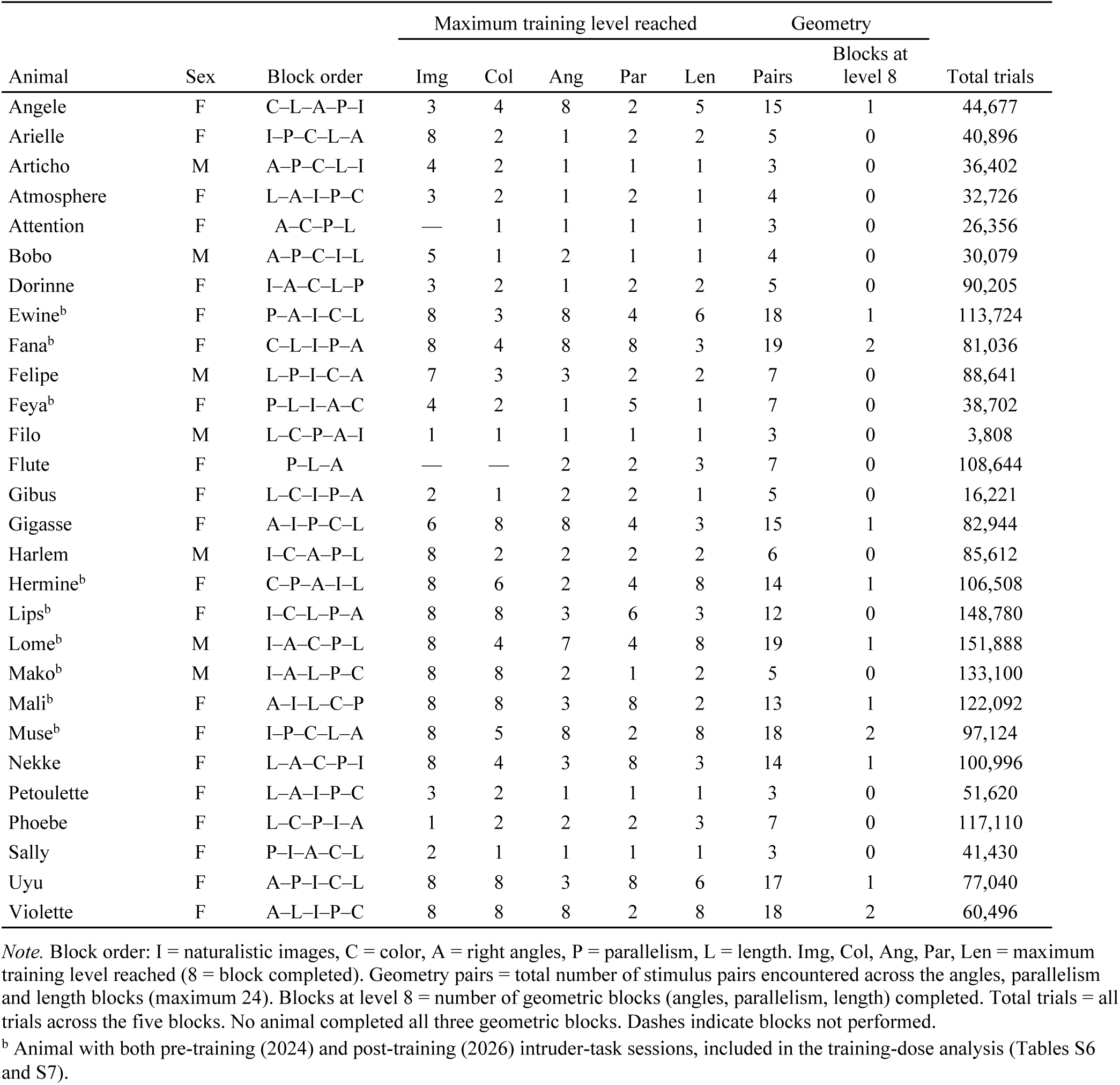
Training history of each baboon. Block order codes: I, naturalistic images; C, color; A, right angles; P, parallelism; L, length. Img, Col, Ang, Par and Len give the maximum training level reached in each block (8 = block completed); dashes indicate blocks not performed. Geometry pairs is the number of stimulus pairs encountered across the three geometric blocks (one new pair per training level; maximum 24). Blocks at level 8 is the number of geometric blocks completed; no animal completed all three. Total trials counts all trials across the five blocks. ^a^Animals tested on the intruder task both before (2024) and after (2026) geometry training, included in the training-dose analysis (Tables S6 and S7).

|  |  |  | Maximum training level reached |  |  |  |  | Geometry |  |
|---|---|---|---|---|---|---|---|---|---|
| Animal | Sex | Block order | Img | Col | Ang | Par | Len | Pairs | Blocks at level 8 | Total trials |
| Angele | F | C–L–A–P–I | 3 | 4 | 8 | 2 | 5 | 15 | 1 | 44,677 |
| Arielle | F | I–P–C–L–A | 8 | 2 | 1 | 2 | 2 | 5 | 0 | 40,896 |
| Articho | M | A–P–C–L–I | 4 | 2 | 1 | 1 | 1 | 3 | 0 | 36,402 |
| Atmosphere | F | L–A–I–P–C | 3 | 2 | 1 | 2 | 1 | 4 | 0 | 32,726 |
| Attention | F | A–C–P–L | — | 1 | 1 | 1 | 1 | 3 | 0 | 26,356 |
| Bobo | M | A–P–C–I–L | 5 | 1 | 2 | 1 | 1 | 4 | 0 | 30,079 |
| Dorinne | F | I–A–C–L–P | 3 | 2 | 1 | 2 | 2 | 5 | 0 | 90,205 |
| Ewine<sup>b</sup> | F | P–A–I–C–L | 8 | 3 | 8 | 4 | 6 | 18 | 1 | 113,724 |
| Fana<sup>b</sup> | F | C–L–I–P–A | 8 | 4 | 8 | 8 | 3 | 19 | 2 | 81,036 |
| Felipe | M | L–P–I–C–A | 7 | 3 | 3 | 2 | 2 | 7 | 0 | 88,641 |
| Feya<sup>b</sup> | F | P–L–I–A–C | 4 | 2 | 1 | 5 | 1 | 7 | 0 | 38,702 |
| Filo | M | L–C–P–A–I | 1 | 1 | 1 | 1 | 1 | 3 | 0 | 3,808 |
| Flute | F | P–L–A | — | — | 2 | 2 | 3 | 7 | 0 | 108,644 |
| Gibus | F | L–C–I–P–A | 2 | 1 | 2 | 2 | 1 | 5 | 0 | 16,221 |
| Gigasse | F | A–I–P–C–L | 6 | 8 | 8 | 4 | 3 | 15 | 1 | 82,944 |
| Harlem | M | I–C–A–P–L | 8 | 2 | 2 | 2 | 2 | 6 | 0 | 85,612 |
| Hermine<sup>b</sup> | F | C–P–A–I–L | 8 | 6 | 2 | 4 | 8 | 14 | 1 | 106,508 |
| Lips<sup>b</sup> | F | I–C–L–P–A | 8 | 8 | 3 | 6 | 3 | 12 | 0 | 148,780 |
| Lome<sup>b</sup> | M | I–A–C–P–L | 8 | 4 | 7 | 4 | 8 | 19 | 1 | 151,888 |
| Mako<sup>b</sup> | M | I–A–L–P–C | 8 | 8 | 2 | 1 | 2 | 5 | 0 | 133,100 |
| Mali<sup>b</sup> | F | A–I–L–C–P | 8 | 8 | 3 | 8 | 2 | 13 | 1 | 122,092 |
| Muse<sup>b</sup> | F | I–P–C–L–A | 8 | 5 | 8 | 2 | 8 | 18 | 2 | 97,124 |
| Nekke | F | L–A–C–P–I | 8 | 4 | 3 | 8 | 3 | 14 | 1 | 100,996 |
| Petoulette | F | L–A–I–P–C | 3 | 2 | 1 | 1 | 1 | 3 | 0 | 51,620 |
| Phoebe | F | L–C–P–I–A | 1 | 2 | 2 | 2 | 3 | 7 | 0 | 117,110 |
| Sally | F | P–I–A–C–L | 2 | 1 | 1 | 1 | 1 | 3 | 0 | 41,430 |
| Uyu | F | A–P–I–C–L | 8 | 8 | 3 | 8 | 6 | 17 | 1 | 77,040 |
| Violette | F | A–L–I–P–C | 8 | 8 | 8 | 2 | 8 | 18 | 2 | 60,496 |

*Note.* Block order: I = naturalistic images, C = color, A = right angles, P = parallelism, L = length. Img, Col, Ang, Par, Len = maximum training level reached (8 = block completed). Geometry pairs = total number of stimulus pairs encountered across the angles, parallelism and length blocks (maximum 24). Blocks at level 8 = number of geometric blocks (angles, parallelism, length) completed. Total trials = all trials across the five blocks. No animal completed all three geometric blocks. Dashes indicate blocks not performed.
<sup>b</sup> Animal with both pre-training (2024) and post-training (2026) intruder-task sessions, included in the training-dose analysis (Tables S6 and S7).

**Table S6.**
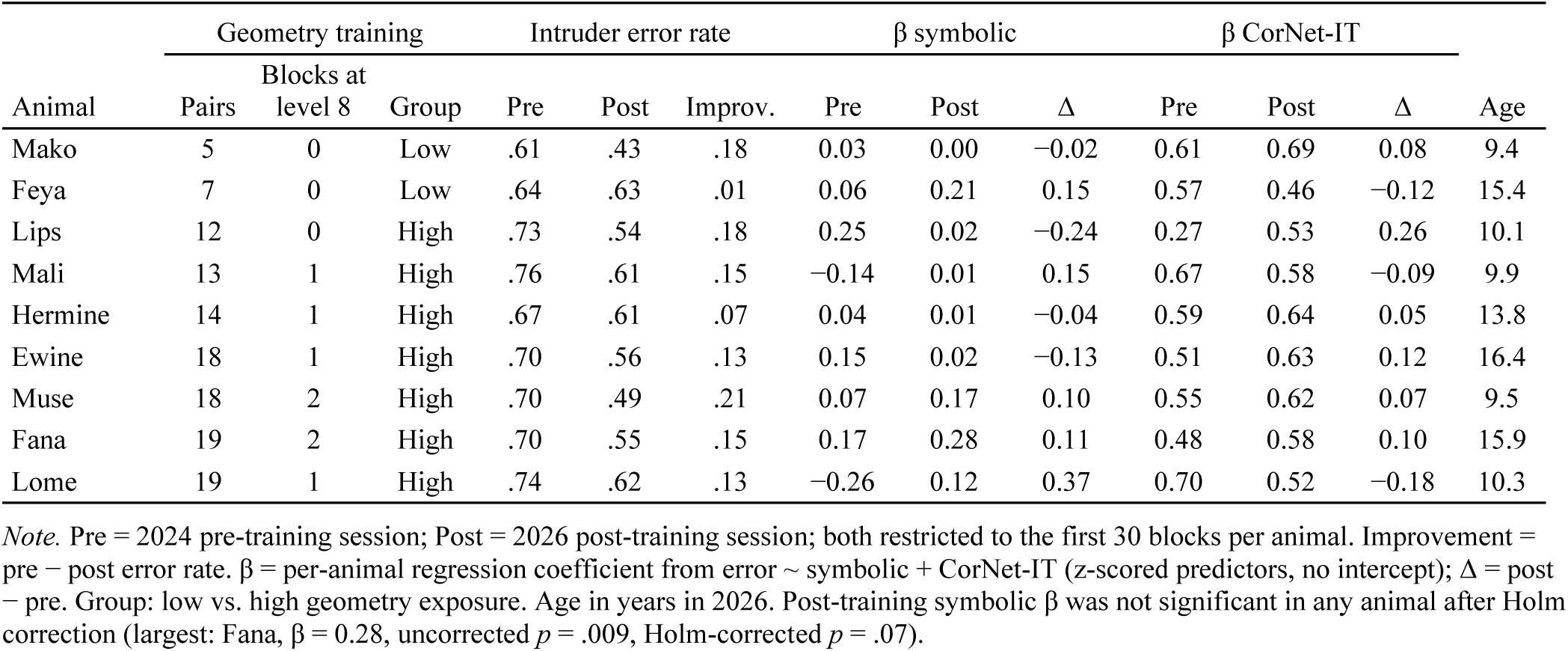
Geometry training dose and change in intruder-task performance for each baboon. Pre is the 2024 session before geometry training and Post the 2026 session after it; both are restricted to each animal’s first 30 blocks. Improvement = pre − post error rate. β symbolic and β CorNet-IT are per-animal coefficients from a regression of intruder-task error on the symbolic-model and CorNet-IT predictors (z-scored, no intercept); Δ = post − pre. Group: low vs. high geometry exposure [≥12 pairs]. Age is in years in 2026. No post-training symbolic β survived Holm correction (largest: Fana, β = 0.28, uncorrected *p* =.009, Holm-corrected *p* =.07).

| Animal | Geometry training |  |  | Intruder error rate |  |  | β symbolic |  |  | β CorNet-IT |  |  | Age |
|---|---|---|---|---|---|---|---|---|---|---|---|---|---|
|  | Pairs | Blocks at level 8 | Group | Pre | Post | Improv. | Pre | Post | Δ | Pre | Post | Δ |  |
| Mako | 5 | 0 | Low | .61 | .43 | .18 | 0.03 | 0.00 | −0.02 | 0.61 | 0.69 | 0.08 | 9.4 |
| Feya | 7 | 0 | Low | .64 | .63 | .01 | 0.06 | 0.21 | 0.15 | 0.57 | 0.46 | −0.12 | 15.4 |
| Lips | 12 | 0 | High | .73 | .54 | .18 | 0.25 | 0.02 | −0.24 | 0.27 | 0.53 | 0.26 | 10.1 |
| Mali | 13 | 1 | High | .76 | .61 | .15 | −0.14 | 0.01 | 0.15 | 0.67 | 0.58 | −0.09 | 9.9 |
| Hermine | 14 | 1 | High | .67 | .61 | .07 | 0.04 | 0.01 | −0.04 | 0.59 | 0.64 | 0.05 | 13.8 |
| Ewine | 18 | 1 | High | .70 | .56 | .13 | 0.15 | 0.02 | −0.13 | 0.51 | 0.63 | 0.12 | 16.4 |
| Muse | 18 | 2 | High | .70 | .49 | .21 | 0.07 | 0.17 | 0.10 | 0.55 | 0.62 | 0.07 | 9.5 |
| Fana | 19 | 2 | High | .70 | .55 | .15 | 0.17 | 0.28 | 0.11 | 0.48 | 0.58 | 0.10 | 15.9 |
| Lome | 19 | 1 | High | .74 | .62 | .13 | −0.26 | 0.12 | 0.37 | 0.70 | 0.52 | −0.18 | 10.3 |

*Note.* Pre = 2024 pre-training session; Post = 2026 post-training session; both restricted to the first 30 blocks per animal. Improvement = pre − post error rate. β = per-animal regression coefficient from error ~ symbolic + CorNet-IT (z-scored predictors, no intercept); Δ = post − pre. Group: low vs. high geometry exposure. Age in years in 2026. Post-training symbolic β was not significant in any animal after Holm correction (largest: Fana, β = 0.28, uncorrected *p* = .009, Holm-corrected *p* = .07).

